# Intercellular mRNA trafficking via membrane nanotubes in mammalian cells

**DOI:** 10.1101/137836

**Authors:** Gal Haimovich, Christopher M. Ecker, Margaret C. Dunagin, Elliot Eggan, Arjun Raj, Jeffrey E. Gerst, Robert H. Singer

## Abstract

RNAs have been shown to undergo transfer between mammalian cells, though the mechanism behind this phenomenon and its overall importance to cell physiology is not well understood. Numerous publications have suggested that RNAs (microRNAs and incomplete mRNAs) undergo transfer via extracellular vesicles (*e.g*. exosomes). However, in contrast to a diffusion-based transfer mechanism, we find that full-length mRNAs undergo direct cell-cell transfer via cytoplasmic extensions, called membrane nanotubes (mNTs), which connect donor and acceptor cells. By employing a simple co-culture experimental model and using single-molecule imaging, we provide quantitative data showing that mRNAs are transferred between cells in contact. Examples of mRNAs that undergo transfer include those encoding GFP, mouse β-actin, and human Cyclin D1, BRCA1, MT2A, and HER2. We show that intercellular mRNA transfer occurs in all co-culture models tested (*e.g.* between primary cells, immortalized cells, and in co-cultures of immortalized human and murine cells). Rapid mRNA transfer is dependent upon actin, but independent of *de novo* protein synthesis, and is modulated by stress conditions and gene expression levels. Hence, this work supports the hypothesis that full-length mRNAs undergo transfer between cells through a refined structural connection. Importantly, unlike the transfer of miRNA or RNA fragments, this process of communication transfers genetic information that could potentially alter the acceptor cell proteome. This phenomenon may prove important for the proper development and functioning of tissues, as well as host-parasite or symbiotic interactions.

**Significance:** Messenger RNA (mRNA) molecules convey genetic information within cells, beginning from genes in the nucleus to ribosomes in the cell body, where they are translated into proteins. Here, we show a novel mode of transferring genetic information from one cell to another. Contrary to previous publications suggesting that mRNAs transfer via extracellular vesicles, we provide visual and quantitative data showing that mRNAs transfer via membrane nanotubes and direct cell-to-cell contact. We predict that this process has a major role in regulating local cellular environments with respect to tissue development and maintenance, cellular responses to stress, interactions with parasites, tissue transplants, and the tumor microenvironment.

**Author contributions:** G.H., A.R. and R.H.S. conceived the research and designed the experiments; C.M.E. performed and analyzed the experiments with WM983b+/-GFP, including transwell and exosomes; M.C.D. and E.E. performed and analyzed the WM983b/NIH393 co-culture experiments; G.H. performed and analyzed all other experiments; and G.H., J.E.G, A.R. and R.H.S. wrote the paper.

## Introduction

An essential aspect of multicellular organisms is the ability of cells to communicate with each other over both long and short distances to coordinate cellular and organ processes. Although most studies have focused on small molecule- or protein-based intercellular communication, little is known about whether RNA molecules act by themselves as mediators of communication. Earlier reports suggested that RNAs may undergo transfer from one cell to another (1–3). However, only recently has it become evident that the extracellular fluids of animals (*e.g.* saliva, plasma, milk, and urine) contain RNAs, including mRNAs (or fragments thereof) and microRNAs (miRNAs). These RNAs are mainly found in extracellular nanovesicles (EVs) such as exosomes (4), although free extracellular ribonucleoprotein (RNP) particles have been identified (5). By using DNA microarrays (4, 6, 7) or RNA-sequencing (RNA-seq) (8), the content of EVs was shown to include a multitude of mRNAs and miRNAs. Thus, it has been proposed that the transfer of RNA between donor and acceptor cells could play an important physiological role.

Yet, examples of exosome-mediated intercellular mRNA transfer and its effects are few (4, 9–12). Only two studies provide qualitative evidence for the possible translation of transferred mRNA in recipient cells (4, 12) and no follow-up research was performed. In particular, there is a lack of quantitative data regarding the number and fate of transferred mRNA molecules at the single cell level. Moreover, the existence of multiple types of EVs (*e.g*. exosomes, microvesicles and apoptotic bodies) that can contain different kinds of cargo, including DNA (13), complicates the study and understanding of this process. In addition, it has been demonstrated that exosomes are transported into lysosomes upon internalization by recipient cells (13–15) and it is unclear how their RNA content reaches sites of translation within the cytoplasm.

Another consideration is that while the abundance of mRNAs in terms of species/copy number in EVs is unknown, the small volume of exosomes might not even allow more than a limited number of mRNA molecules per exosome. The presence of full-length, functional mRNAs in exosomes was demonstrated in a few studies (4, 12), whereas others suggest that exosomes contain mainly RNA fragments (16, 17). Indeed, one study found that only short mRNA fragments (<500 nucleotides [nts]) are efficiently loaded into EVs, as compared to longer transcripts (>1500nts) (16). Recent reports also suggest that miRNAs are present in very low abundance in EVs (*i.e.* individual miRNAs may range between 10^-4^ to 60 miRNA molecules/exosome in a given extracellular fluid) (18, 19), suggesting that miRNA transfer via EVs might have little influence on recipient cells unless selective mechanisms for uptake exist. Overall, these results suggest that a large number of EVs containing specific mRNAs (or mRNA fragments) or miRNAs might be necessary in order to significantly affect the physiology of recipient cells. Additionally, other modes of mRNA transfer were not investigated. Thus, an unbiased quantitative approach is needed to determine if mRNA is indeed transferrable, how is it transferred, and how much is transferred.

To study intercellular mRNA transfer in a quantitative and unbiased manner, we employed a simple strategy, which is depicted in Fig. 1A-B. In this model, “donor” and “acceptor” cells are co-cultured together, and the transfer of specific mRNA species from donors to acceptors is visualized and quantified by single-molecule Fluorescent *In Situ* Hybridization (smFISH) (20, 21) or live imaging using the MS2 aptamer system (22). Donor and acceptor pairings can consist of cell types from any typical mammalian species (*e.g.* rat, mouse, human), provided that the query mRNA is expressed only in the donor cells. By using this model, we discovered that mRNAs can transfer between cells and provide absolute quantitative data on the number of transferred mRNA molecules per cell under different culture conditions.

**Figure 1.**
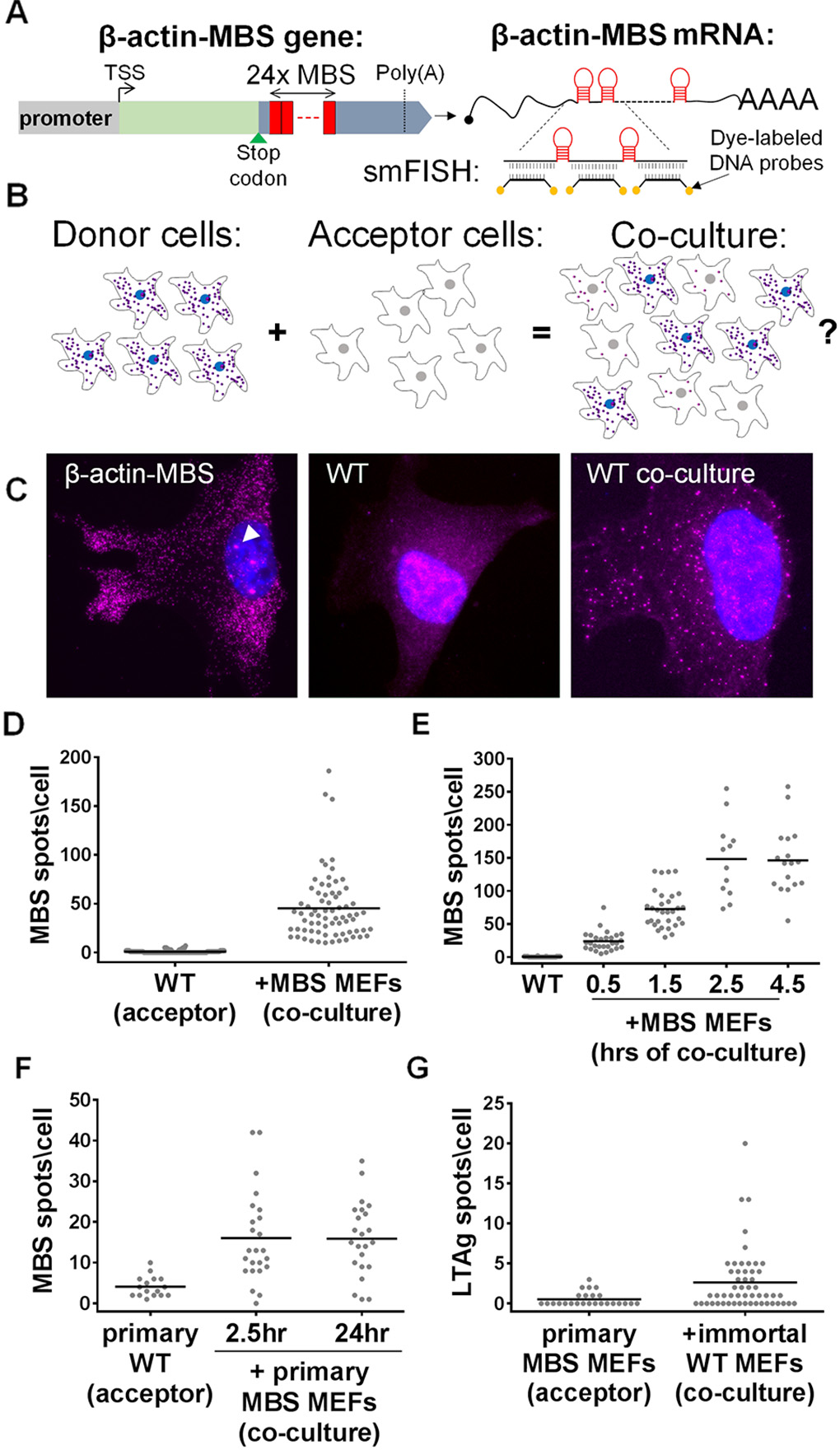
Detection of β-actin-MBS mRNA transfer by smFISH. **(A)** A schematic depicting the β-actin-MBS gene (left) and resulting mRNA (right). mRNA detection was accomplished using smFISH with fluorescence-labeled DNA probes against the MBS sequence (aka MBS probes). *TSS* – transcription start site. *Poly(A)* – polyadenylation site. **(B)** A schematic illustrating the basic experimental set-up. Donor cells (left) that express a unique mRNA (*e.g*. β-actin-MBS; shown as small purple dots) were co-cultured with naïve acceptor cells (middle) that lack this mRNA. If mRNAs undergo transfer from donor to acceptor cells, co-culture yields labeling of the acceptor cells (right). Each cell type was also cultured separately to assess mRNA expression levels (in donor cells) or background staining (in acceptor cells). **(C)** smFISH images of an immortalized donor MBS MEF, immortalized acceptor WT MEF, and acceptor WT MEF in co-culture. Labels: blue - DAPI staining of the nucleus; Magenta - Cy3-tagged MBS probes. Arrowhead indicates a transcription site. **(D)** Distribution of the number of β-actin-MBS mRNA spots observed in immortalized WT MEFs cultured alone or co-cultured with donor MBS MEFs for 24hrs. Each dot in this and all other panels (*E-G*) represents the score of the number of mRNAs detected in a single cell as obtained by smFISH using MBS probes. Bar (in panels *D-G*): mean number of spots per acceptor cell. **(E)** Distribution of the number of β-actin-MBS mRNA spots in WT MEFs as a function of time after co-culture with donor MBS MEFs. **(F)** Distribution of the number of β-actin-MBS mRNA spots in primary WT MEFs (pWT) cultured alone or co-cultured for 2.5 or 24hrs with primary donor MBS MEFs. **(G)** Distribution of the number of LTag mRNA spots in primary MBS (pMBS) MEFs co-cultured with LTag-immortalized donor WT MEFs for 24hrs. See Table S1 for data on *N* (number of cells scored), mean, standard error of the mean (SEM), and *p*-values for each experiment.

We show that mRNA transfer requires direct cell-to-cell contact and that it occurs via membrane nanotubes (mNTs; also known as tunneling nanotubes) and not by diffusion. mNTs are long and thin cytoplasmic projections involved in direct contact-dependent intercellular communication between eukaryotic cells. mNTs were shown to support cell-to-cell transfer of small molecules, proteins, prions, viral particles, vesicles and organelles in a variety of cell types (23–33). Here we demonstrate that mNTs can also transfer mRNA molecules and identify mRNAs encoding a wide variety of proteins that undergo intercellular transfer in *in vitro* culture conditions.

## Results

### mRNA can transfer between cells

To determine whether cell-cell mRNA transfer occurs, immortalized wild-type (WT) mouse embryonic fibroblasts (MEFs) were co-cultured with immortalized MEFs derived from a homozygous transgenic mouse that harbors 24 repeats of the MS2-coat protein (MCP) binding sequence (MBS) at the 3’ untranslated region (UTR) of the endogenous alleles of β-actin (referred to here as MBS MEFs) (22). smFISH with MBS-specific probes was used to analyze the number of β-actin-MBS mRNAs detected and quantitation was performed using in-lab programs or FISH-quant (34) (see SI Materials and Methods). MBS MEFs showed up to several thousand distinct FISH spots in each cell, as well as bright nuclear foci representing transcription sites (Figs. 1C, left panel and S1, and Table S1) (22). Immortalized MBS MEFs are tetraploid and have up to four transcription sites (22). As expected, β-actin-MBS mRNAs and transcription sites were not detected in WT MEFs cultured alone (Fig. 1C, middle panel). However, when co-cultured with MBS MEFs for 24hrs, WT cells acquired MBS-labeled mRNAs (Fig. 1C, right panel) at an average (± standard error of mean; SEM) of 45± 4 mRNAs/cell, and as many as ~190 mRNAs/cell (Fig. 1D and Table S1).

To determine the global rate of mRNA transfer, we measured the number of transferred β-actin-MBS mRNAs in WT MEFs at 0.5, 1.5, 2.5 and 4.5hrs after adding MBS MEFs to the culture. Under these conditions, MBS MEFs attached to the fibronectin-coated glass surface within 15-20min. We detected transferred mRNA within 30min of co-culture (*i.e.* 10-15min after MBS MEFs attached to the surface). The number of transferred mRNAs increased with time until reaching a plateau at 2.5hrs after co-culture (Fig. 1E and Table S1).

Zipcode-binding protein 1 (ZBP1) is an RNA-binding protein (RBP) previously shown to be required for β-actin mRNA localization to the leading edge and focal adhesions in fibroblasts (35, 36) and to dendrites in neurons (37, 38). However, the absence of ZBP1 in the donor MBS MEFs (*i.e.* immortalized β-actin-MBS zbp1^-/-^ MEFs) did not hinder mRNA transfer to immortalized acceptor WT MEFs (Fig. S2A and Table S1).

To determine that mRNA transfer is not due to immortalization, we examined whether it occurs between primary cells. Primary MEFs derived from WT or MBS mice were co-cultured for either 2.5 or 24hrs and smFISH was performed to detect β-actin-MBS mRNA transfer. Similar to immortalized MEFs, transferred β-actin-MBS mRNA was detected in co-cultured primary WT MEFs (Fig. 1F and Table S1). This indicated that intercellular RNA transfer is not unique to immortalized cells. Co-cultures of primary MEFs and immortalized MEFs yielded a higher level of mRNA transfer (*i.e.* 2-fold) when compared to primary co-culture (Fig. S2B and Table S1). Co-culturing primary and immortalized MEFs also allowed us to test the transfer of a second mRNA, SV40 large T antigen (LTag) mRNA, which is expressed only in the immortalized cells (see Fig. S3A and B, and Table S1 for expression levels in donor cells). By employing LTag-specific smFISH probes, we could detect the transfer of LTag mRNA from immortalized to primary MEFs (Fig. 1G and Table S1). This indicates that transfer is not unique to β - actin mRNA or to MBS-labeled mRNAs.

FISH experiments using Cy3-labeled MBS- and Cy5-labeled open reading frame (ORF)-specific probes showed that an average of 3.5± 0.4% of the total β-actin mRNA found in WT MEFs (as detected by ORF-specific probes) was transferred from donor MBS cells (Fig. S4A and B, and Table S1). In these FISH experiments, most of the MBS spots detected in MBS MEFs were co-localized with ORF spots, although there were a few single-color labeled spots (Fig. S4C). It is important to note that many MBS spots detected in WT MEFs also co-localized with ORF spots (Fig. S4D and E) indicating that the transferred β-actin-MBS mRNAs detected in acceptor cells constituted full-length transcripts and not solely 3’UTR or MBS fragments.

### mRNA transfer occurs in heterologous human/murine cell co-cultures

To test the generality of this process, we first determined if β-actin-MBS mRNA from MEFs can transfer to other cell types, including human cells. We therefore co-cultured MBS MEFs with a human embryonic kidney cell line (HEK293T) and examined these cells for β-actin-MBS mRNA. Indeed, we found that β - actin-MBS mRNA can transfer from murine to human cells (Fig. 2A and Table S1). Although the mean amount of endogenous β-actin mRNA levels in MEFs is about 3-fold higher than in HEK293T cells (Fig. S4F and Table S1), the transferred mRNA constitutes 4.5± 0.6% of the β-actin ORF spots in HEK293T cells (Fig. S4G and H, and Table S1). This is similar in percentage to the amount of transferred mRNA between MEFs, which suggests that this is either a regulated or limited process. β-actin-MBS mRNA transfer was also detected in co-cultures of MBS MEFs with the human osteosarcoma (U2OS) or adenocarcinoma (SKBR3) cell lines (Fig. S5A and B, and Table S1). This shows that the mechanism of transfer is conserved and confers the murine-human exchange of mRNA. Reciprocal transfer experiments using human-specific probes showed that an endogenous mRNA, such as BRCA1 mRNA (Fig. S3A and B, and Table S1), transferred from HEK293T or HEK293 cells to MEFs (Fig. 2B and Table S1). Likewise, endogenously expressed SERP2, MITF, and MT2A mRNAs (Fig. S3A and B, and Table S1) transferred from human melanoma cells (WM983b-GFP) to murine embryonic fibroblasts (NIH3T3) in co-culture (Fig. 2C-E and Table S1). The transfer of the ectopically expressed GFP mRNA (Fig. S3A and B, and Table S1) could also be detected both in human-murine and human-human co-cultures (Fig. S5C-D and Table S1). For these experiments, we employed smFISH probes that tiled the entire length of the transcript for the non-MBS labeled mRNAs. In some cases (*e.g.* GFP, SERP2, MITF and MT2A), dual-color probe sets (39) were used to ascertain the specific identification of these mRNAs, since two-color co-localization enhances the probability that the signal is specific (Fig. S5D). These approaches strongly indicate that full-length mRNAs underwent transfer. To test for transfer of a different type of RNA molecule, we also examined by dual-color smFISH whether a highly expressed (*e.g.* >2000 copies/cell) human-specific long non-coding RNA (lncRNA), MALAT1 (39), underwent transfer, but observed the transfer of only 1-2 molecules in a small percentage (~8%) of acceptor cells (Fig. S5F and Table S1). At the moment we cannot determine whether the lack of appreciable MALAT1 RNA transfer is due to its localization in the nucleus (39) or its specific function and/or regulation.

**Figure 2.**
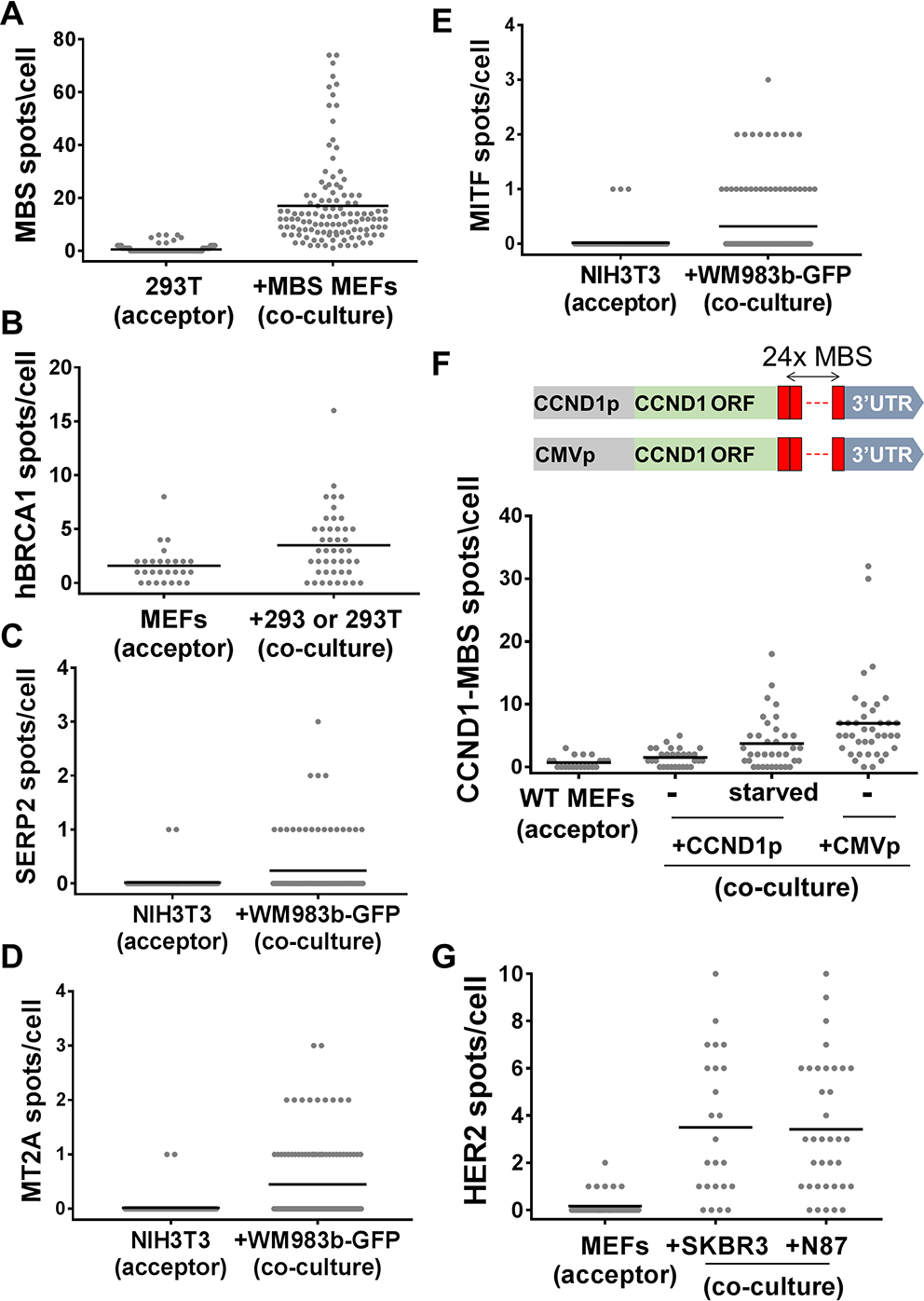
Transfer of mRNAs in human-murine co-cultures. **(A)** Distribution of the number of β-actin-MBS mRNA spots in HEK293T cells that were co-cultured with donor MBS MEFs for 24hrs. Each dot in this and all other panels (*B-G*) represents the score of a single cell using smFISH. Bar (in panels *A-G*): mean number of spots per acceptor cell. **(B)** Distribution of the number of endogenously expressed human BRCA1 mRNA spots in WT MEFs co-cultured for 24hrs with donor HEK293T or HEK293 cells, as detected using sequence-specific probes against BRCA1. **(C-E)** Distribution of the number of SERP2 (C), MITF (D) and MT2A (E) mRNA spots in NIH3T3 cells that were co-cultured with donor WM983b-GFP human melanoma cells for 48hrs, as detected using sequence-specific probes. Only dual-color spots were considered as legitimate mRNA spots. **(F)** Distribution of the number of Cyclin D1-MBS mRNA spots in WT MEFs co-cultured for 7hrs with donor HEK293 cells expressing Cyclin D1-MBS mRNA from the endogenous promoter (CCND1p) or the CMV promoter (CMVp). “Starved” - WT MEFs were serum starved over-night prior to co-culture. MBS probes were used for the detection of CCND1-MBS mRNA. The diagram above the plot illustrates the CCND1-MBS gene under the two different promoters in the HEK293 cell lines. **(G)** Distribution of the number of endogenously expressed human HER2 mRNA spots in MEFs co-cultured for 3hrs with donor SKBR3 or NCI-N87 cells, using sequence-specific probes. See Table S1 for data on *N*, mean, SEM and *p*-values.

### Gene expression in donor cells may influence mRNA transfer

In contrast to β-actin-MBS mRNA, which undergoes transfer at tens to hundreds of molecules per cell (Figs. 1C-F, 2A, S2A and B, S4B and H, and S5A and B), only a small amount of the other RNAs tested were transferred (Figs. 1G, 2B-E, and S5C-D). Furthermore, primary MBS MEFs, which express less β - actin-MBS mRNA than immortalized MBS MEFs (Fig. S1), also exhibited less transfer (Fig. S2B). Thus, we hypothesized that mRNA expression levels in donor cells might affect the absolute number of transferred RNAs.

To test whether gene expression affects mRNA transfer, two HEK293 cell lines that express MBS-labeled Cyclin D1 (CCND1-MBS) mRNA from either its endogenous promoter (CCND1p) or a CMV promoter (CMVp) were obtained (40). As expected, CMVp induced higher levels of expression of CCND1-MBS mRNA in donor cells compared to CCND1p (Fig. S3A and B, and Table S1). We co-cultured WT MEFs with either of these cell lines and compared the level of CCND1-MBS mRNA transfer. In agreement with our hypothesis, more RNA transfer was detected when the MEFs were co-cultured with HEK293 cells bearing CMVp-CCND1-MBS (Fig 2F and Table S1).

To further explore this issue, we examined the transfer of HER2 mRNA from human cell lines to MEFs. We used two epithelial cell lines having different expression levels of HER2: gastric carcinoma cells (NCI-N87; 316± 22 mRNAs/cell) and adenocarcinoma cells (SKBR3; 611± 36 mRNAs/cell) (Fig. S3A and B, and Table S1). We observed mRNA transfer in both cases and despite the elevated expression of HER2 mRNA in SKBR3 cells we observed the same low level of transfer (*e.g.* ~ 3.5 mRNAs/cell; Fig. 2G and Table S1). Given that HER2 mRNA is highly expressed in the donor cells, similar to β-actin, this result indicates that factors other than gene expression level may influence transfer.

### Stress conditions affect mRNA transfer

To determine whether intercellular mRNA transfer is affected by external physiological conditions, we examined the effect of stress upon mRNA transfer. In these experiments, either donor or acceptor cells were exposed to stress (*e.g*. heat-shock, oxidative stress, protein-folding stress, or serum starvation) prior to co-culture. Cells were relieved from the stress and the reciprocal cells (*i.e.* unstressed acceptor or donor cells) were plated on top. Under these conditions, recovery from heat-shock inhibited mRNA transfer, whereas recovery from oxidative stress (H_2_O_2_ treatment), protein-folding stress (DTT treatment), or serum starvation increased the extent of mRNA transfer (Figs. 2G and 3, and Table S1). Interestingly, stress conditions modulated mRNA transfer mostly when applied to the acceptor cells. Thus, different stresses may make cells either less or more receptive to mRNA transfer, but have little effect on the capacity of donor cells to transfer mRNA once the stress is relieved. This occurs even though β-actin-MBS mRNA levels increased in the stressed donor cells after they were allowed to recover (Fig. S1). This also implies that mRNA expression levels alone may not be the only parameter that influences transfer. We next tested whether translational stress inhibits mRNA transfer. We found that β-actin-MBS mRNA transfer was not inhibited by cycloheximide (Fig. S6 and Table S1), a translation inhibitor, indicating that this process does not require *de novo* protein synthesis.

### mRNA transfer requires direct cell-to-cell contact

The isolation and characterization of exosomes and other EVs from different cell types has revealed that these vesicles may contain miRNAs and mRNA (or at least mRNA fragments) and, therefore, could serve as a means of mRNA transfer between cells (4, 12). Thus, we speculated that intercellular mRNA transfer is mediated by EVs that convey their contents by diffusion through the medium and uptake into acceptor cells. To test this hypothesis, we transferred “conditioned” media from donor cell cultures (*e.g.* MBS MEFs or WM983b-GFP cells) to acceptor cell cultures (*e.g.* WT MEFs or WM983b cells, respectively) and looked for transferred β-actin-MBS or GFP mRNA in the acceptor cells following 1.5-2.5 or 24hrs of incubation. Transferred mRNA molecules were not detected in the acceptor cells (Fig. 4A and B, and Table S1), indicating that the mode of transfer is not via the growth medium. Consequently, to determine if mRNA transfer is mediated by physical contact, donor and acceptor cells were co-cultured under conditions that do not allow for contact between cells, but allow for the sharing of diffusible materials. We used two different approaches - the first approach, which we term “tripod”, is illustrated in Fig 4C. In this system, donor and acceptor cell layers are physically separated by several millimeters, though any type of particle can diffuse across the medium. The second approach, termed “transwell”, utilized a physical barrier to separate the cell layers and exclude the transfer of particles >0.4-5µm (Fig. 4D). In either case, little to no evidence for mRNA transfer was detected and only under co-culture conditions that allow for physical contact was transfer observed (Fig. 4A and B, and Table S1). Lastly, exosomes were directly isolated from WM983b-GFP cells. These exosomes were 40-60nm in diameter (Fig. S6B) and contained primarily small RNAs (<200nt) (Fig. SC). The isolated exosomes were applied to WM983b GFP-negative cells. After 24hrs of incubation with the acceptor cells no appreciable mRNA transfer was detected (Fig. 4B and Table S1). Thus, mRNA transfer appears to require cell-cell contact.

Although the release of mRNAs from dying cells might also allow for transfer, we observed little cell death in our MEF co-culture experiments (~3%). Nevertheless, we tested whether cell death contributes to transfer by pretreating donor MBS MEFs with H_2_O_2_ (3%, 1.5 hrs) to induce oxidative stress and apoptosis. This treatment resulted in ~45% cell death during the subsequent 2.5hr of incubation in co-cultures with acceptor MEFs using the tripod approach described above. We observed only a slight increase in transferred β-actin-MBS mRNA levels in acceptor MEFs (Fig. 4A and Table S1). This level of transfer is far less than that observed under co-culture conditions using healthy cells, indicating that the release of apoptotic bodies cannot account for mRNA transfer under normal growth conditions. In parallel experiments we used an apoptosis-inducing drug, raptinal (41), to selectively kill donor cells during co-culture. Raptinal leads to apoptosis within <2hrs by inducing cytochrome c release from mitochondria, which in turn activates the apoptotic cascade. Consistent with published results (41), APAF-1 knock-out (APAF-1^k/o^) MEFs are resistant to short-term (1hr) treatment with the drug, whereas APAF-1^+/+^ cells (*e.g.* MBS MEFs) were highly sensitive and showed >95% cell death within <2hr. We next co-cultured MBS MEFs and APAF-1^k/o^ MEFS for 3 or 12hrs and then treated the cells with raptinal for 1hr. Transferred mRNA was detected by smFISH after co-culture but before treatment (*i.e.* time “0”) and at different time points during drug treatment (*e.g.* 30min, 1hr) or after drug wash-out (*e.g.* 2-6hrs). Importantly, the amount of transferred β-actin-MBS mRNA was reduced to very low levels upon raptinal treatment (Fig. S7A and Table S1). Some MBS MEF cell fragments containing mRNA were detected between cells, suggesting that β-actin-MBS mRNA can be present and remain intact in apoptotic bodies (Fig. S7B, panel i). Likewise, we observed clusters of transferred mRNAs in a few acceptor cells (Fig S7B, panel ii) suggesting that apoptotic bodies were engulfed by these cells. Nevertheless, the contribution of apoptotic bodies to mRNA transfer appears to be extremely limited and does not contribute to the mechanism observed in healthy cells (Fig. S7A). Thus, mRNA transfer in co-cultures is not mediated by apoptotic bodies.

### mRNA transfer occurs via membrane nanotubes

To test if mRNA transfer requires close proximity or direct contact, cells were plated at a 99:1 ratio of WT to MBS MEFs and co-cultured for 24hrs. smFISH was performed to determine the amount of β-actin MBS mRNA transferred to either nearest or distant neighbor cells. Nearest neighbors were defined as those directly proximal to the MBS cells (*i.e.* residing in the adjacent cell layer), while distal cells constitute those in the surrounding 2^nd^ or 3^rd^ layers. We observed that the number of MBS spots in adjacent cells was higher than in cells located further away (Fig. S8A-B and Table S1).

Our results indicate that a proximity-based mechanism confers intercellular mRNA transfer. To determine whether mRNAs are transferred directly via known cell-cell contacts (*e.g.* gap junctions), we treated MBS and WT MEF co-cultures with 100μM carbenoxolone, a gap junction inhibitor (42), for 60-90min. However, we found no effect of carbenoxolone upon β-actin-MBS mRNA transfer (Fig. S8C and Table S1). By eliminating other possibilities (*e.g.* diffusion or gap junctions), we suspected that mRNA transfer might occur via membrane nanotubes (mNTs). mNTs are long (up to ~200µm), thin (0.05-0.5μm), cellular protrusions that can transfer many types of components from one cell to another (23–33). Indeed, upon examination of our co-culture images, we could detect the presence of mRNAs in mNT-like structures (Figs. 4E-F and S9A). These were fairly rare images, since the visibility of mNTs (as detected by the background fluorescence of the FISH protocol) was weak. Furthermore, we suspect that many mNTs are destroyed during the FISH process. Indeed, we could more easily detect mNTs by live imaging (see below).

To further substantiate the role of mNTs in mRNA transfer, we employed known mNT inhibitors. It was previously shown that fibronectin (FN)-coated glass supports mNT formation better than polylysine (PL)-coated glass (43). Consistent with this finding, we found that cells plated on uncoated or PL-coated glass exhibited much less mRNA transfer when compared to cells plated onto FN-coated glass (Fig. S9B and Table S1). An alternative explanation for the reduction of transfer on PL-coated glass was if the expression levels of mRNA in the donor cells were greatly reduced. Yet, upon examination the expression levels of β-actin-MBS mRNA in the donor MBS MEFs were only ~30% less under these conditions (Fig. S1). Next, we tested the effects of the actin depolymerization drug, Latrunculin A (LatA), and the CDC42 inhibitor, CASIN, which were previously shown to inhibit mNT formation or the transfer of proteins through mNTs (24, 44). Consistent with the involvement of mNTs, we found that treatment of WT/MBS MEF co-cultures with either LatA or CASIN resulted in a two-fold reduction in β-actin mRNA transfer (Fig. S9C-D and Table S1). The reduced level of transfer cannot be attributed to decreased mRNA levels in donor cells (Fig. S1) and, furthermore, the percentage of transferred mRNA present in LatA-treated co-cultures was greatly reduced (Fig. S9E and Table S1). Thus, mRNA transfer through contact-dependent mNTs seems to be the likely mechanism.

### Live imaging of mRNA transfer

A great advantage of the MS2 labeling system is the ability to follow mRNA movement in real time by live imaging. Although we could detect transferred β-actin-MBS mRNA in acceptor cells after co-culture using both smFISH with probes against MBS and immunofluorescence (IF) using anti-GFP antibodies to detect tandem MCP-GFP (tdMCP-GFP) (45), the number of co-labeled spots was very low (Fig. S10A). Indeed, the expression of tdMCP-GFP in the donor cells appeared to greatly reduce the level of transfer in co-cultures (Fig. S10B and Table S1). In contrast, the expression of either tdMCP-GFP in the acceptor cells or GFP alone in the donor cells did not lower the level of mRNA transfer. Thus, the binding of tdMCP-GFP to β-actin-MBS mRNA in the donor cells appears to inhibit mNT-mediated delivery of mRNA to acceptor cells. While it was difficult to detect transfer by live cell imaging, nevertheless, we documented one clear event of linear β-actin-MBS mRNA transfer between donor and acceptor cells (Fig. 5A-B and Movie S1). The rate of β-actin-MBS mRNA movement in Movie S1 was calculated at 4.85µm/min. This rate is similar to that of other components that transfer via mNTs (46). In addition, we often observed mNTs by live imaging and on rare occasions detected mRNAs moving along the length of mNT-like structures or appearing in acceptor cells using tdMCP-GFP (Figs. 5C-D and S9Aiii, and Movies S2-6).

### Discussion

Current research suggests that cells secrete RNA molecules into extracellular fluids, which are then taken up by downstream acceptor cells to alter gene expression and, ultimately, cell physiology. Although the evidence for miRNA transfer via EVs or RNP particles is compelling, the evidence for EV-mediated transfer of mRNA is lacking, both in qualitative and quantitative terms. Here, we took an unbiased approach to ask whether intact mRNA molecules are transferred between cells. We provide visual evidence and quantitative data showing that mRNA molecules undergo intercellular transfer and that this transfer occurs via mNTs between adjacent cells and not by diffusion (see model, Figure 6). This work presents the results of independent studies performed and validated by different research teams.

#### Do all mRNAs transfer?

The data presented in this study show that essentially all mRNAs tested can undergo transfer between mammalian cells (Figs. 1–2 and S5). This list includes native endogenously-expressed mRNAs (*e.g.* β-actin-MBS, MITF, SERP2, MT2A, BRCA1, and HER2), ectopically-expressed mRNAs (*e.g.* GFP, LTag, CCND1-MBS), as well as MS2 aptamer-tagged mRNAs. These different mRNAs share no known sequence commonalities, nor do their encoded proteins localize and/or function on the same cellular processes or pathways. Moreover, the list includes both non-mammalian (GFP) and viral (LTag) proteins. Overall, the results suggest that perhaps all mRNAs are amenable to transfer. Thus far, the only exception we have identified is MALAT1, a lncRNA that resides primarily in the nucleus. However, it is unclear whether the lack of MALAT1 transfer is due to its localization or because it is a non-coding RNA. Since use of smFISH has limited our analysis to only a small number of genes, non-biased genome-wide transfer experiments that necessitate high-throughput approaches, such as RNA-seq or MERFISH (47), are needed to allow for the detection of large numbers of individual transcripts. This will allow us to define and quantitate the extent of the RNA transferome via the large-scale identification of transferrable versus non-transferable mRNAs and lncRNAs. Such an approach may help identify *cis* elements or epi-transcriptomic changes that recruit proteins involved in RNA transfer. This may allow us to predict which RBPs associate with transferred mRNAs and thereby facilitate the transfer process. Importantly, our FISH-IF experiment indicates that the tdMCP-GFP protein is not removed from β-actin-MBS mRNA upon transfer (Fig. S10). Thus, cellular proteins involved in transfer might remain bound to the transferred mRNA in acceptor cells and pulldown of these mRNAs could reveal the identity of these RBPs. Another aspect of selectivity is how specific mRNA molecules are chosen for transfer (*i.e.* out of the total pool for any given mRNA species)? While also unclear, our results suggest that translation does not play a role, since translation inhibition did not affect β-actin-MBS mRNA transfer (Fig. S6).

#### Is mRNA transfer solely expression-dependent?

Two single-cell approaches are used to quantify the number of mRNA transcripts in cells: single-cell RNA sequencing (scRNA-seq) and smFISH. The detection level of scRNA-seq depends upon the methods of single cell isolation, RNA extraction and depth of sequencing, but may suffer from amplification biases and transcript underestimation when compared to spike-in controls (48). Thus, the sensitivity of scRNA-seq may not provide accurate detection when there are <10 mRNA molecules/cell (48). In contrast, mRNA visualization by smFISH allows for unbiased measurements at single molecule resolution, while maintaining the integrity of cell structure and conferring spatial resolution of the detected mRNA molecules. By using smFISH as our method of choice, we detected low numbers of transferred mRNA molecules in acceptor cells; the average for many being <10 molecules/cell. In contrast, β-actin-MBS mRNA was exceptional in that hundreds of mRNA molecules/cell could undergo transfer in co-culture experiments.

What makes β-actin-MBS mRNA so effective at transfer? One possible explanation is that high levels of β-actin-MBS mRNA expression in donor cells increases the likelihood for transfer. The idea that mRNA transfer correlates with gene expression is further supported by the finding that elevation of CCND1-MBS mRNA, using the CMV promoter, increased the number of transferred molecules (Fig. 2F). However, high gene expression levels alone may not guarantee higher levels of transfer. For example, we observed similar levels of HER2 mRNA transfer from two different donor lines that had very different levels of expression (Figs. 2G and S3). Likewise, the expression of MITF, MT2A and SERP mRNAs were differentially expressed in the same donor cells, but led to an equally low level of transfer (Figs. 2C-E and S3). We do note, however, that low levels of transfer might be due to the fact that the cells were plated on uncoated glass in these experiments. This reduces the efficiency of mRNA transfer, in comparison to cells plated on fibronectin-coated glass (Fig. S9B). That said, the transfer of CCND1-MBS, LTag, BRCA1, and HER2 mRNAs was also relatively low in comparison to β-actin-MBS mRNA, and these cells were plated on fibronectin-coated glass. Therefore, we propose that gene expression is probably but one factor that determines mRNA transfer.

Aside from gene expression, other factors influence the propensity of a given mRNA to be transferred. These include cell culture conditions (*e.g*. Fig. S9B), acceptor cell stress (Fig. 3), cell type specificity (*e.g.* N87 cells vs. SKBR3; Figs. 2G and S3), sequence elements or epi-transcriptomic modifications, and RBPs specific to the mRNA in question. Other factors may also be considered - for example, promoter elements are known to affect the cytoplasmic fate of mRNAs (49). Hence, it is possible that elements at the CMV promoter are responsible for the elevated rate of CCND1-MBS mRNA transfer, rather than its elevated expression *per se* (Figs. 2F and S3). Organellar localization of an mRNA (*e.g.* nuclear retention) could also affect availability. Furthermore, it is possible that mRNAs involved in mNT formation (*e.g.* β-actin) or spatially distributed near mNTs might show a greater propensity for transfer. This issue will have to be resolved in future studies.

**Figure 3.**
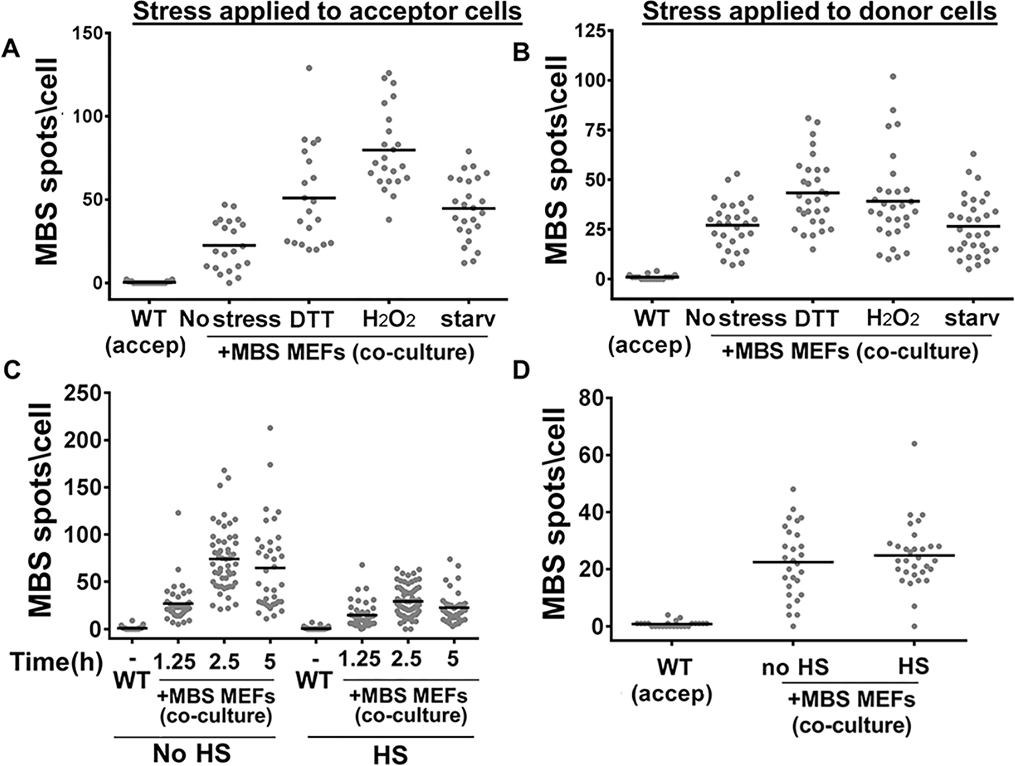
Stress affects β-actin-MBS mRNA transfer. **(A)** Acceptor WT MEFs or **(B)** donor β-actin-MBS MEFs cells were either left untreated (No stress) or treated for 1.5hrs with dithiothreitol (DTT) (1mM), H_2_O_2_ (1mM), or serum starvation (Starv), prior to scoring for mRNA transfer by smFISH with MBS probes. **(C)** Acceptor WT MEFs or **(D)** donor β-actin-MBS MEFs cells were exposed to heat-shock (HS, 42°C) for 1hr or were left untreated (No HS). Following stress, the reciprocal cell line was plated and co-culture was maintained under stress-free conditions for 2.5hrs **(A, B, D)** or for the indicated times **(C).** β-actin-MBS mRNA transfer was detected by smFISH using MBS probes, as described in Fig 1. See also Table S1 for data on N, mean, SEM and *p*-values. Bar (in panels *A-D*): mean number of MBS spots per acceptor cell.

#### MCP-GFP inhibits mRNA transfer

The MS2 system has been widely used to image mRNAs within many organisms and cell types (50), and has not been shown to have deleterious effects upon mRNA movement. Furthermore, a mouse model that expresses both β-actin-MBS and MCP-GFP in all cells did not show physiological, developmental, or behavioral defects (51). Yet, we found that the expression of tdMCP-GFP in donor MBS MEFs inhibited the transfer of β-actin-MBS mRNA (Fig. S10B) and Cyclin D1-MBS mRNA (not shown). The reduction in RNA transfer cannot be explained by a reduced expression level of β-actin-MBS mRNA, since tdMCP-GFP did not affect steady-state levels (Fig. S1). Time-lapse imaging of hundreds of live cells for varying durations and at various intervals between frames led to only a single clear example of mRNA transfer (Fig. 5a-B and Movie S1), and only a few examples of mRNAs residing in mNTs (Figs. 5C-D and S9Aiii and Movies S2-4,6). This might explain why transfer was not detected earlier in other studies employing MS2-labeled mRNAs for single-molecule imaging.

While the cause of tdMCP-GFP-mediated inhibition of β-actin-MBS mRNA transfer is not known, we presume that the larger size/mass of the mRNP particle impedes interactions with the transport machinery and/or recruitment into mNTs. Formation of this complex may invariably slow the anterograde movement of mRNA through mNTs in comparison to retrograde transport, leading to a net movement back to the donor cells (Movie S6). Another possibility is that tdMCP-GFP binding to the mRNA results in structural changes in the RNA that interfere with the binding of factors essential for transfer.

Clearly, inhibition of β-actin mRNA transfer between cells is not deleterious at the organismal level, since β-actin-MBS x MCP-GFP crosses are fully viable (51). This is probably because β-actin is ubiquitously expressed in all cell types and, therefore, is not expected to be limiting. We predict, however, that the inhibition of transfer of other mRNA species might yield more obvious and deleterious effects at both the cellular and organismal levels. The loss of transfer of cell type-specific mRNAs may alter the physiology of downstream acceptor cells, although this will have to be determined on a case-by-case basis.

#### What is the mechanism of mRNA transfer?

Our work demonstrates that mRNAs transfer between cells via mNTs and not via diffusion-based mechanisms. Yet, the mechanisms that regulate nanotube formation and maintenance are not well understood. Likewise, even less is known how mRNAs are recruited to mNTs and undergo trafficking therein.

mNTs are thought to be actin filament-based, yet microtubules may also be present and possibly exist as the sole cytoskeletal structure (52). In our study, the inhibition of actin polymerization was found to reduce β-actin-MBS mRNA transfer in co-cultured cells (Fig. S9D), which implies at least a partial role for actin. The CDC42 small GTPase, which regulates actin filament formation, has also been implicated in mNT formation (24, 53), perhaps in the elongation rather than initiation phase (54). Nevertheless, the inhibition of CDC42 reduced mRNA transfer (Fig. S9C), strengthening our hypothesis that mRNAs transfer via actin-based mNTs.

Other proteins have been shown to modulate mNT formation. For example, TNFaip2/M-Sec utilizes the RalA GTPase and exocyst complex to initiate mNT formation (53, 54), and are recruited along with filamin and myosin to the plasma membrane by the MHC class III protein, LST1 (44), to initiate mNT formation. Although it is yet unclear which cytoskeletal motors are responsible for mNT-mediated mRNA transfer, the velocity of the RNP particle shown in Movie S1 strongly resembles that of myosin motors (55, 56). In particular, Myosin Va is involved in RNA trafficking (57) and is probably a good candidate to explore given that earlier work suggested a role for this motor in the distribution of Schwann cell-synthesized RNA to neuronal cell bodies and axons after lesioning (58). Preliminary analyses performed by us using myosin-inhibiting drugs did not yield conclusive results revealing which myosin motor is involved (data not shown). However, future experiments employing the knock-down/knock-out of specific myosin motors should elucidate which motor confers mNT-mediated mRNA transfer.

Finally, the question arises of whether mRNAs are transferred in free RNP particles or in particles bound to cellular membranes? Organelles, such as intact mitochondria, as well as lysosomes, endosomes, Golgi, and endoplasmic reticulum (ER) have been shown to transfer via mNTs (26, 29, 59, 60). Likewise, mRNAs are known to associate with and undergo intracellular co-trafficking with organelles (*e.g.* ER, mitochondria, and peroxisomes; (61–64). Thus, we speculate that organelle-localized mRNAs may undergo transfer along with the organelle. Live imaging of mNT-mediated mRNA transfer using cells expressing labeled organelles should help resolve this issue.

#### Why do mRNAs transfer between cells?

The biological importance of mRNA transfer between cells is still unknown. Clearly, mRNAs or their fragments are found in EVs and are presumably taken up by the surrounding cell layers in tissues. Our discovery of mNT-mediated mRNA transfer suggests that full-length mRNAs can also be exchanged between cells through contact. Although we cannot rule out that this phenomenon occurs only under *in vitro* culture conditions, it would seem unlikely given the existence of mNTs in tissues and their known ability to transfer intracellular material. Thus, we presume that the process of intercellular mRNA transfer is active (*i.e.* cytoskeleton- and motor-dependent), is responsive to environmental cues (*e.g*. stress), and effects downstream cellular responses.

mNTs have been characterized primarily in cell and tissue cultures. However, they were also detected in patient-derived solid tumors (65). This suggests a role for mNTs in cancer biology and the tumor microenvironment. The finding that primary MEFs can be mRNA donors, as well as acceptors (Figs. 1F-G and S2B), indicates that mRNA transfer is not a consequence of immortalization or tumorigenesis *per se*. Therefore, this process is expected to occur in embryonic and/or normal adult tissues and may affect development, maintenance, or both. Future studies employing human xenografts in rodents may help to resolve this question.

Presuming that mNT-mediated mRNA transfer occurs in animals, the main biological question is what impact the transfer of a few mRNA molecules has upon downstream acceptor cells? The answer depends on the mRNA in question and hinges on whether the transferred mRNA is translated and at what efficiency. For instance, cancer cells that transfer a few mRNA molecules encoding a key transcription factor not normally expressed in untransformed neighbor cells might induce or repress the transcription of genes that regulate responses to extracellular signals elicited from the cancerous cells and, thereby, facilitate cancer cell motility or the supportive nature of tumor local microenvironment. It is possible to speculate that mRNAs with transforming potential (*i.e.* oncogenes) could induce carcinogenesis in neighboring cells upon transfer. Likewise, the transfer of mRNAs involved in cell differentiation during embryonic development might act as means to induce or repress neighboring cells. Determining the scope of this novel process and deciphering the mechanism and physiological outcome of mRNA transfer will be the goal of future studies.

### Materials and Methods

#### Cells and cell lines

Primary mouse embryonic fibroblasts (MEFs) from WT (C57BL/6) or β-actin-MBS mice were isolated from E14.5 embryos and cultured as is or immortalized by transfection with SV40 large T antigen, as previously described (22). Immortalized ZBP1 knock-out (ZBP1^-/-^)-MBS MEFs were described earlier (36). HEK293T, U2OS, NIH3T3 and SKBR3 cells were purchased from ATCC. These cell lines were received as gifts: SKBR3 (M. Oren, Weizmann Institute of Science - WIS); U2OS (Z. Livneh, WIS); N87 (Y. Yarden, WIS); WM983b (M. Herlyn, The Wistar Institute); HEK293 cells expressing P_CCND1_-CCND1-MBS or P_CMV_-CCND1-MBS(40) (Y. Shav-Tal, Bar Ilan University); and SV40-immortalized APAF-1 knock-out MEFs and isogenic wild-type MEFs(66) (M. Orzáez, Centro de Investigación Príncipe Felipe (CIPF), Spain).

MEFs expressing EGFP, NLS-HA-tdMCP-GFP (referred to here as tdMCP-GFP) or palmitoylated TagRFP-T (TagRFP-T-ps), were created by infection with the appropriate lentivirus, followed by sorting by flow-cytometry to isolate only infected cells. Cells were sorted for low expression levels of EGFP, NLS-HA-tdMCP-GFP, and for high levels of expression of TagRFP-T-ps. WM983b-GFP cells were created by clonal selection, as previously described (67).

#### Plasmids and lentivirus generation

A lentivirus vector (pHAGE-UBC-RIG) carrying NLS-HA-tdMCP-GFP was previously described (45) (Addgene plasmid #40649). DNA sequences encoding EGFP and TagRFP-T-ps were cloned into the same viral backbone vector. Plasma membrane (inner leaflet)-associated palmitoylated TagRFP-T (TagRFP-T-ps) was generated by the addition of a sequence encoding 20 amino acids of rat GAP-43 (MLCCMRRTKQVEKNDEDQKI)(68) to the 5’ end of the TagRFP-T gene.

Lentivirus particles were produced by transfecting the expression vector along with plasmids for ENV (pMD2.VSVG), packaging (pMDLg/pRRE) and REV (pRSV-Rev) (Addgene plasmids #12259, #12251 & #12253, respectively) into HEK293T cells using calcium phosphate(69). The virus-containing supernatant was harvested and concentrated using a Lenti-X concentrator (Clontech), per the manufacturer’s instructions. Virus particles were resuspended in Dulbecco’s modified Eagle’s medium (DMEM) containing 10% Fetal Bovine Serum (FBS), aliquoted, and stored at -80°C for subsequent infection of cells in culture.

#### Cell culture conditions

MEFs, HEK293, HEK293T and U2OS cells were cultured routinely in 10cm dishes in DMEM (4.5gm/l glucose) supplemented with 10% FBS, 1mM sodium pyruvate, and antibiotics (0.1mg/ml streptomycin and 10U/ml penicillin) at 37°C with 5% CO_2_. Primary MEFs were cultured in the same medium at 37°C with 10% CO_2_ and 3% O_2_. SKBR3 and N87 cells were cultured in RPMI1640 medium supplemented with 10% FBS, 1mM sodium pyruvate, and antibiotics. RPMI1640 medium was also used for the co-culture of either SKBR3 or N87 cells with MEFs that were pre-conditioned to RPMI1640. WM983b cells were cultured in Tu 2% medium (78.4% MCDB153 medium, 19.6% Leibovitz’s L-15 medium, 2% FBS, and 1.68mM CaCl_2_).

Fibronectin (10µg/ml in PBS, Sigma) was used to coat glass coverslips (round 18mm #1) for FISH experiments and the glass-bottom dishes (MatTek Cat #: P35G-1.5-14-C) for live imaging. For some experiments, poly-D-lysine (Sigma) was used to coat coverslips at 1mg/ml or 0.1mg/ml, as indicated. Cells were dissociated from the dishes using 0.25% trypsin-EDTA and plated on freshly coated coverslips. We found that the co-culturing of both donor and acceptor cells immediately after dissociation tends to reduce the transfer efficiency of β-actin-MBS mRNA (data not shown). Therefore, acceptor cells were typically plated the day before co-culture (*i.e.* in the afternoon/evening) and donor cells then plated on top the next morning, unless otherwise indicated. The acceptor:donor ratio was 1:1 unless otherwise indicate. Co-culture was performed for 30min-24hrs, as indicated, prior to fixation and FISH analysis as detailed below. For live imaging, cells were cultured on fibronectin-coated glass-bottom dishes, as described above, using DMEM/10%FBS medium. The co-culture was maintained for 1-2hrs in the incubator. During that time, the microscope’s environmental control chamber was warmed to 37°C with humidity control and normal atmosphere. The medium was then replaced with pre-warmed Leibovitz’s L-15 medium lacking phenol red and containing 10% FBS and the cells were taken for imaging. Imaging sessions of live cells lasted between 1-10hrs. Experiments using WM983b cells were performed using uncoated glass and co-cultures (plated at a ratio of 1:1) were incubated for 48hrs prior to FISH analysis. In all cases, cell density (upon plating in co-culture) was calculated to achieve ~80± 10% confluence at the time of fixation or live imaging.

For “tripod” experiments, paraffin was heated to ~110°C. By using a glass pipette, paraffin drops (2-3mm in height) were placed at three points on the coverslip edges (*i.e.* in a triangular arrangement). Once the paraffin solidified, coverslips were exposed to UV light (using the tissue culture hood lamp) for 30min. Tripods were stored under sterile conditions at room temperature until used. For transwell experiments, Corning Transwell™ multiple-well plates with permeable polycarbonate 0.4µm or 5µm membrane inserts (Fisher Scientific Cat. # 07-200-147 or 07-200-149) were used.

The drugs cycloheximide (CHX, 100µg/ml; Sigma), 2-[(2,3,4,9-Tetrahydro-6-phenyl-1*H*-carbazol-1-yl)amino]ethanol (CASIN, 1µM; gift of V. Krizhanovsky, WIS), Latrunculin A (LatA, 200nM; Santa Cruz Biotechnology), carbenoxolone (100µM; Sigma) and raptinal (10µM; gift of P. Hergenrother, U. Illinois Urbana-Champaign) were added directly to the media. Drugs were added 20-30min after plating of the donor cells (*i.e.* after MBS MEFs have attached). For the induction of protein folding or oxidative stress, dithiothreitol (DTT, 0.1mM; Sigma) or H_2_O_2_ (1mM) was added directly to the media and cells were further incubated for 1.5hrs. Serum starvation was induced by replacing the media with DMEM lacking FBS and the cells further incubated for the indicated times. Heat-shock was induced by submerging cells that were pre-cultured on coverslips in a sealed 12-well plate for 1hr in a 42°C water bath. Following incubation under stress conditions, the stressed cells were washed with pre-warmed media, upon which the other (non-stressed) cell type was added, and the cells were co-cultured for the indicated times under stress-free conditions. For the apoptosis-tripod experiment, cells grown on “tripod” coverslips were first treated with 3mM H_2_O_2_ for 1.5hrs prior to co-culture. After an additional 2.5hrs of culture in media lacking H_2_O_2_ ~45% of MBS MEFs were dead. The percentage of cell death was determined using trypan blue staining.

#### Exosomes purification

In order to isolate exosomes, media were collected from two cultures of WM983b-GFP cells in T75 flasks (70-90% confluence, 48hrs culture) and exosome isolation was performed, as previously described (70). Briefly, cells were cultured in medium supplemented with exosome-free serum (*i.e.* serum depleted of exosomes was first generated by ultracentrifugation of the serum overnight at 120,000 x *g* at 4°C followed by decanting of the liquid phase). Media were collected after 48hrs of cell culture and centrifuged at 300 x *g* for 10min at 4°C to remove cells and debris. The supernatant was centrifuged at 16,500 x *g* for 20min at 4°C and the resulting supernatant filtered through a 0.2µm membrane filter (SteriFlip) and then ultracentrifuged at 120,000 x *g* for 70min at 4°C to pellet the exosomes. The exosome pellet was resuspended in 250µl of phosphate-buffered saline (PBS). Alternatively, exosomes were isolated using Total Exosome Isolation Reagent^™^ (ThermoFisher, Cat #: 4478359). Particle size of isolated exosomes was measured using a Zetasizer instrument (Malvern). RNA extracted from whole cells and isolated exosomes (miRNeasy, Qiagen) was analyzed using a Bioanalyzer 2100 (Agilent). Exosomes were applied directly onto WM983b cells cultured on a two-well glass bottom plate (LabTek Cat#: 155379). Cells were incubated with the exosomes for 24hrs prior to FISH.

#### Fluorescent *in situ* hybridization (FISH) and FISH-immunofluorescence (FISH-IF)

Tiled FISH probes (20-mers) against the MBS sequence (comprising three oligos with amino-allyl moieties on both the 5’ and 3’ ends) and β-actin ORF (comprising 35 5’ and 3’ amino-allyl oligos) (22) were end-labeled with Cy3 or Cy5 (GE Healthcare), as previously described (71). Tiled FISH probes (20-mers) against human BRCA1-Quasar (Q)570 (ShipReady Cat#: SMF-2028-1), human HER2-Q570 (DesignReady Cat# VSMF-2102-5), and custom probe sets against LTag-Q670 mRNAs were obtained from Biosearch Technologies (Petaluma, CA, USA). Tiled odds/evens dual-color 20-mer probes against GFP and human MITF, SERP2, MT2A and MALAT1 were purchased as 3’-amine oligos from Biosearch Technologies. These oligos were coupled to Cy3 or Alexa Fluor 594 (Life Technologies) fluorophores and purified by HPLC, as previously described (39).

FISH was performed at the Singer/Gerst labs as previously described (22), with slight modifications (see a detailed protocol in SI Materials and Methods). FISH at the Raj lab was performed on WM983b cells according to Biosearch Stellaris^®^ RNA FISH protocol.

For FISH-IF experiments, cell fixation and permeabilization were performed as described for FISH. Pre-hybridization was performed in PHB supplemented with 3% BSA (Sigma) and RNase inhibitor (10U/ml Superase (Ambion) or RNasin (Promega)). For hybridization, 20U/ml Superase or RNasin and primary chicken (IgY) anti-GFP antibody (GFP-1010, Aves labs; 1:5000), as well as the FISH probes, were added to the hybridization mix. Samples were incubated in humid chamber in the dark at 37°C for exactly 3hrs. Following hybridization, coverslips were rinsed twice in PHB, then incubated twice for 30min at 37°C in PHB supplemented with 3% BSA and secondary goat anti-chicken IgY antibody conjugated with Alexa Fluor 647 (Life Technologies A21449; 1:1000). Samples were further washed, DAPI stained and mounted on slides as for FISH.

#### Imaging

FISH and FISH-IF images were taken using different microscopes, as detailed in SI Materials and Methods. Exposure times for imaging varied between different cell cultures, probes, dyes and microscopes, and were determined empirically per experiment. All slides from the same experiment were imaged using the same illumination parameters. Live imaging was performed at AECOM on an Olympus IX-71 Total Internal Reflection Fluorescence (TIRF) station customized for laser-illumination as detailed in SI Materials and Methods.

#### Image analysis and data presentation

Microscope images presented in the figures and movies were minimally processed for brightness and contrast using FIJI program (72). The plots depicted in Fig. S2E were generated in FIJI by using the “strait line” tool to measure pixel intensity.

Analysis of smFISH images to quantify FISH spots was performed either using in-house developed MATLAB programs or FISH-quant (FQ) (34). Airlocalize (22) was used at the Singer lab for the experiments depicted in Figs. 1C,E, 4A and S8B. All experiments involving WM983b cells were analyzed at Raj lab with Rajlabimagetools (73). FQ was used in the analysis of all other experiments. For more details, see SI Materials and Methods. The data presented in all graphs and in Table S1 represents data collected from two or more experiments. In a few cases that showed distinct sub-populations (*i.e*. Figs. S2A and B, and S7A) the different experiments were color-coded. The data in Fig. S1 (immortalized) was pooled from all experiments with MBS MEFs.

**Figure 4.**
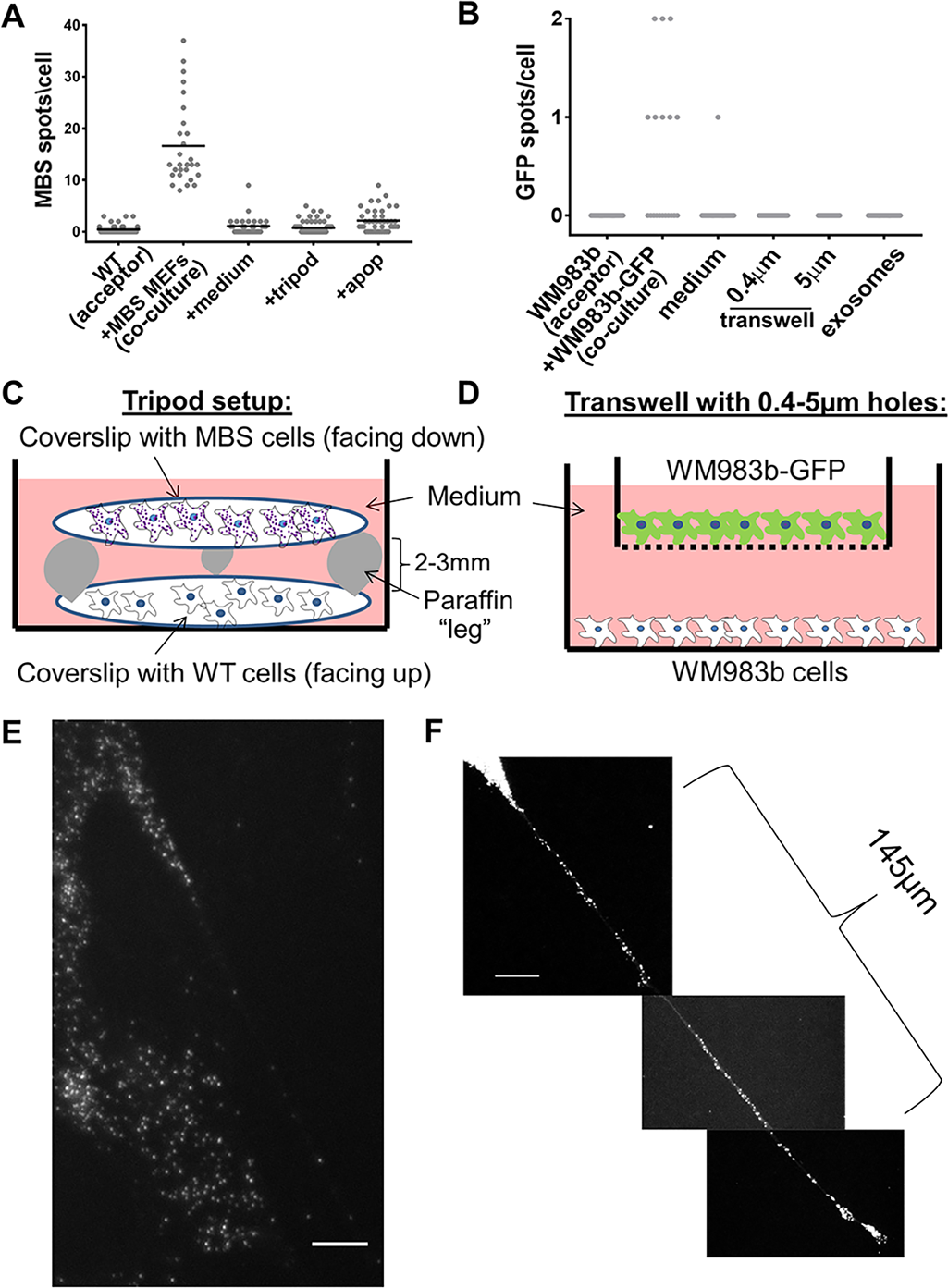
Intercellular mRNA transfer requires direct cell-to-cell contact. **(A)** Distribution of the number of β-actin-MBS mRNA spots in acceptor WT MEFs co-cultured with donor of β-actin-MBS cells, as shown in Fig. 1 (co-culture), as incubated with media collected from an overnight culture of MBS MEFs (+medium), as shown in the tripod setup (+tripod), or as shown in the tripod setup with dying cells (+apop). Apoptosis was induced in donor MBS cells by pre-treatment with 3% H_2_O_2_ prior to co-culture in the tripod set-up with WT MEFs. Incubation time was 2hrs for each treatment Bar: mean number of MBS spots per acceptor cell. **(B)** Distribution of the number of GFP mRNA spots in acceptor WM983b cells co-cultured with donor WM983b-GFP cells (co-culture). WM983b cells were incubated either with media collected from an overnight culture of WM983b-GFP cells (medium) in the transwell setup (using either 0.4~m or 5~m pores) or with exosomes isolated from WM983b-GFP cells. Incubation time was 24hrs for each treatment. Detection was performed using GFP-specific probes **(C)** A schematic depicting the tripod setup. Donor and acceptor cells grown separately on glass coverslips were positioned facing each other, but separated by 2-3mm using paraffin legs. **(D)** A scheme depicting the transwell setup. WM983b-GFP cells were cultured in the upper chamber of transwells of different porosity before being transferred to transwells containing WM983b cells plated on the bottom chamber **(E)** smFISH image of β-actin-MBS mRNA present in a mNT formed by a primary β-actin-MBS MEF. Scale bar: 5µm. **(F)** smFISH image of β-actin-MBS mRNA along a mNT formed by an immortalized β-actin-MBS MEF. Scale bar: 10µm. See Table S1 for data on *N*, mean, SEM and *p*-values.

**Figure 5.**
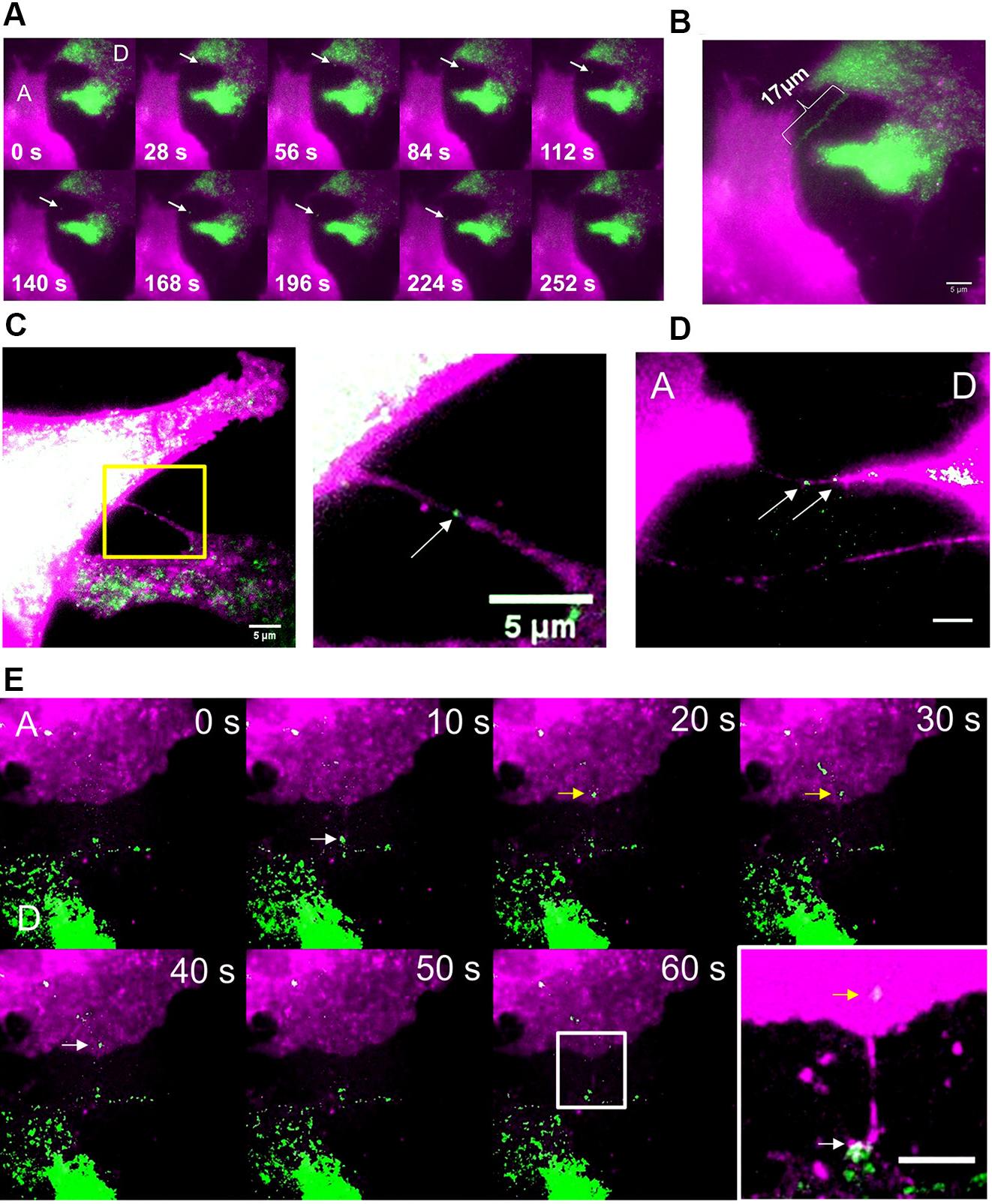
Live imaging of mRNA transfer. **(A)** Individual time-lapse images from Movie S1 of live imaging of β-actin-MBS mRNA (green) transferring from a donor MBS MEF cell (“D”) to an acceptor WT MEF cell (“A”). The membranes of both cells were labeled with TagRFPT (magenta). Arrow points to an mRNA being transferred. **(B)** Maximum projection of Movie S1. Scale bar: 5µm. **(C)** Live image of a β-actin-MBS mRNA in a nanotube connecting donor MBS MEFs. Image taken from a co-culture of WT acceptor MEFs and donor MBS MEFs expressing tdMCP-GFP showing a mNT that connects two MBS cells. Both cells are labeled with membrane targeted-TagRFP-T (magenta) and the arrow points to a tdMCP-GFP-labeled mRNA (green). Image is taken from Movie S2. Scale bar = 5μm. **(D)** Live image of β-actin-MBS mRNAs in a mNT connecting donor and acceptor MEFs. A still image from Movie S4 of cells cultured in *A* shows a mNT that connects a donor MBS MEF (“D”) with an acceptor (“A”) WT MEF. Both cell types are labeled with membrane targeted-TagRFPT (magenta) and the arrows point to tdMCP-GFP-labeled mRNAs (green). Scale bar = 5μm. **(E)** A time-lapse photomontage that may capture β-actin-MBS mRNA transferring from a donor cell to an acceptor. WT acceptor MEFs (“A”) and donor MBS MEFs expressing tdMCP-GFP (“D”) from *A* were co-cultured and monitored for mRNA transfer. Images taken from Movie S5. Arrows points to a tdMCP-GFP-labeled mRNA (green) that appears to transfer between frames (*i.e.* labeled white in the donor cell and yellow in the acceptor cell). The donor cell is denoted by the green label, due to tdMCP-GFP expression, but also shows less TagRFP-T label in this case. The last image is a magnified section (square) of a maximum projection of the movie, showing a nanotube-like structure connecting the two cells at the presumed path of the transferring mRNA.

**Figure 6.**
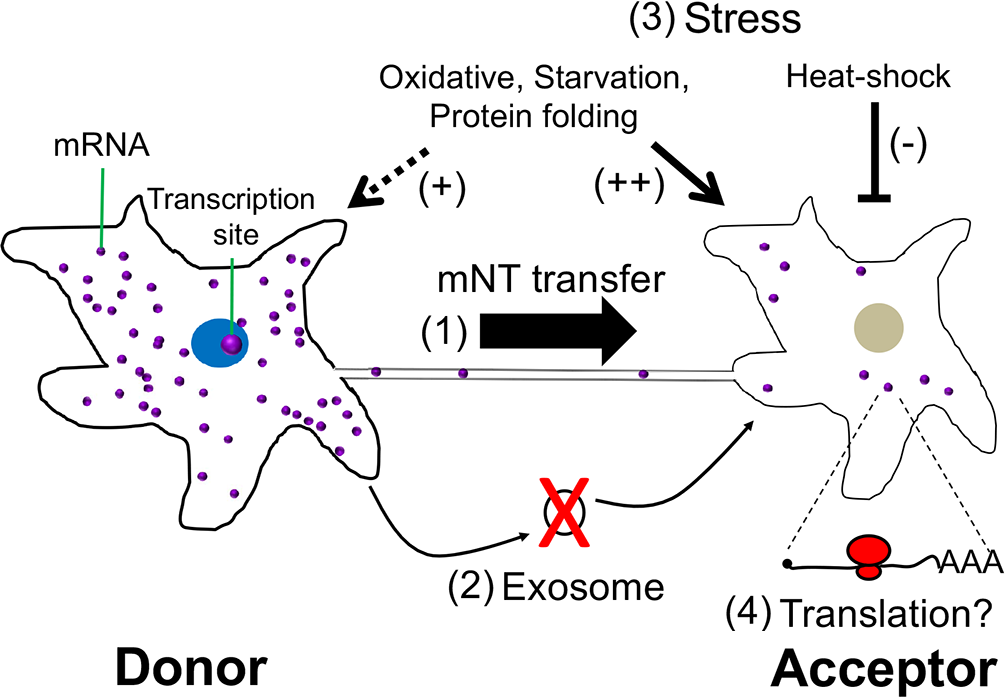
A schematic representation of intercellular mRNA transfer. A donor cell (left) that expresses a given mRNA can transfer mRNA molecules to an acceptor cell (right) through a mNT (1). This process appears to be actin-myosin regulated, requires direct physical contact, and does not occur via exosomes (2). Stress conditions appear to affect mRNA transfer, whereby oxidative, protein-folding, and nutrient stresses on both donor and acceptor cells favor transfer, whereas heat-shock of acceptor cells inhibits transfer (3). Transferred mRNAs are likely to undergo translation, although this has not yet been proven (4).

#### Statistical analysis

Unpaired t tests were used to calculate *p*-values (depicted in Table S1) for each two sets of compared results. All calculations of average, SEM and *p*-values were performed using GraphPad Prism software (GraphPad software Inc).

## Acknowledgments

We thank Xiuhua Meng and Bin Wu for plasmids, Yaron Shav-Tal, Moshe Oren, Yosef Yarden, Meenhard Herlyn and Maria Orzáez for cell lines; Paul Hergenrother for raptinal; Valery Krizhanovsky for CASIN; Shalev Itzkovitz for use of his microscopes, and Florian Mueller for excellent technical support for FISH-quant. G.H. is a recipient of the Gruss Lipper Family Post-doctoral fellowship (EGL Charitable Foundation); the Dean of Faculty (Weizmann Institute of Science) and Sir Charles Clore Post-doctoral fellowships; and the Koshland Foundation and McDonald-Leapman Grant Senior Post-doctoral fellowships. This research was funded by grants from the Joel and Mady Dukler Fund for Cancer Research (WIS) to J.E.G.; US-Israel Binational Science Foundation-National Science Foundation (#2015846) to J.E.G. and R.H.S.; NIH New Innovator 1DP2OD008514 to A.R.; and NIH grant NS083085 to R.H.S.

## Supporting Information - Haimovich et al. (2017)

### Material and Methods

#### Fluorescent *in situ* hybridization (FISH)

Cells cultured on coverslips in 12-well plates were washed three times with PBS supplemented with 5mM MgCl_2_ (PBSM) and then fixed for 10min with 4% paraformaldehyde in PBSM at room temperature (RT). Cells were washed once in PBSM+0.1M glycine for 10min and then twice for 10min in PBSM. In time-course experiments (Figs. 1C, 3C and S7), samples taken at early time points were maintained at 4°C until the collection of samples from all the time points was performed. Cells were permeabilized with 0.1% Triton X-100 (Surfact-Amps X-100, ThermoScientific) in PBS for 10min followed by three 10min washes in PBSM. Cells were incubated for 30min with pre-hybridization buffer (PHB; 10% formamide in 2x saline sodium citrate buffer (2xSSC)). Coverslips were placed, cells facing down, on a 45µl cushion of hybridization buffer [2xSSC, 10% dextran sulfate (Sigma), 10% formamide (Sigma), 1mg/ml *E. coli* tRNA (Roche), 0.2mg/ml BSA (Roche), 2mM vanadyl ribonucleoside complex (VRC, Sigma), and 10ng/sample of labeled FISH probes] on Parafilm^™^ in a humid chamber (*e.g.* a sealed tissue culture dish). Samples were incubated in the dark at 37°C for 3hrs-overnight. Following hybridization, coverslips were returned to 12-well plates, cell side facing up, and incubated twice for 15min in 1ml of PHB at 37°C, rinsed three time with 2xSSC, stained with 4’,6-diamidino-2-phenylindole (DAPI, 0.5µg/ml in 2xSSC) for 1min, and washed for 5min with 2xSSC. Coverslips were mounted on slides using Prolong^™^ gold mounting medium (Molecular Probes) and were left to dry for at least 2hrs prior to imaging (samples were typically imaged after 12-72hrs). Slides were sealed with nail polish (Electron Microscopy Sciences) prior to imaging and were stored at -20°C

#### Microscopes used for Imaging

At AECOM an Olympus BX-61 microscope equipped with an X-cite QH120 PC lamp (EXFO), PlanApo 100× 1.35 NA oil immersion objective (Olympus) and a CoolSNAP ZCCD camera (Photometrics) was used. 0.2µm step *z*-stack images were taken with a MS 2000 XY t WIS,Aautomated stage (ASI) and using MetaMorph software (Molecular Devices) at 0.0645µm/pixel. lympusOthree different microscopes were employed: 1) A Delta Vision (DV) system that employed an nIX-71 microscope equipped with a mercury lamp (Olympus), PlanApo 100× 1.35 NA oil immersio weresobjective (Olympus), and a CoolSNAP HQ CCD camera (Photometrics). 0.2µm step *z*-stack image 2) A taken with an automated stage (Applied Precision) and using DV software at 0.0645µm/pixel; 1.45, aANikon Ti-E inverted fluorescence microscope equipped with a ×100 oil-immersion objective N 024 CCD1wide-spectrum light source (either Prior Lumen 220Pro or Nikon Intensilight), and a Pixis agetcamera (Photometrics). 0.2µm step *z*-stack images were taken with a MS 2000 XYZ automated s (ASI) and using MetaMorph software (Molecular Devices) at 0.130µm/pixel; and 3) A Zeiss thAxioObserver Z1 DuoLink dual camera imaging system equipped with Illuminator HXP 120 V lig source, PlanApo 100× 1.4 NA oil immersion objective and Hamamatsu Flash 4 sCMOS cameras. 0.2µm step *z*-stack images were taken using a motorized XYZ scanning stage 130x100 PIEZO, and ZEN2 software at 0.0645µm/pixel or 0.130µm/pixel. At U. Penn. a Nikon Ti-E inverted Prior fluorescence microscope equipped with a ×100 oil-immersion objective NA 1.4, equipped with Lumen 220Pro and a Pixis 1024BR CCD camera was used. 0.3µm step *z*-stack images were taken. with a Prior automated stage and using MetaMorph software at 0.130µm/pixel Live imaging was performed at AECOM on an Olympus IX-71 Total Internal Reflection 640nm, equipped Fluorescence (TIRF) station customized for laser-illumination at 488nm, 561nm and with an Olympus 60× 1.45 NA TIRFM or Olympus 150× 1.45 NA TIRFM objectives, an environmental control chamber and an Andor EMCCD camera. 0.6µm step *z*-stack images were taken with a MS 2000 XYZ automated stage (ASI) and using MetaMorph software. Imaging was performed in wide-field mode and not in TIRF mode. The duration of imaging, as well as the time interval between frames, number of *z*-sections, exposure times, and laser power were determined empirically for each movie presented (see SI Movie Legends for details).

#### Image analysis using Fish-quant, Airlocalize, and Rajlabimagetools

Spot characterization in Fish-quant (FQ) was initially performed on donor cells and later applied to acceptor cells. Briefly, after background subtraction (“filtration”; typical parameters for filtering in FQ were: Kernel BGD XY, Z = 6, 5; Kernel SNR XY, Z = 0.5, 1), the FISH spots were fitted to a 3D Gaussian. The intensity and the width of the 3D Gaussian were thresholded to exclude non-specific signals. These values were determined empirically for each experiment or imaging session (*i.e.* in cases where the same sample was imaged under different microscopes/conditions). These parameters varied somewhat between cells, probes, dyes, microscopes and experiments. Negative controls (*i.e.* acceptor cells that did not express the tested mRNA) were included in all FISH experiments in order to facilitate thresholding of FISH analysis parameters. The only case for which a negative control was unavailable was that of the β-actin ORF. In this case, the threshold was determined so that the number of β-actin ORF spots in MBS MEFs is similar to MBS spots in these same cells.

Due to the high background seen with FISH probes in the nuclei of cells (particularly in those of MEF acceptor cells), nuclei were excluded from the spot analysis in most cases. Since FQ determines the nuclear outline based on the 2D max-projection image of the DAPI signal, FISH spots that were in *z*-sections above or below the nucleus were excluded from the analysis. After initial parameter settings and thresholding based on a donor cell, all images were analyzed using the batch analysis tool. Thresholding was optimized after batch analysis and images were re-analyzed based on the optimized thresholds. Parameters were set to give as close to zero positive spots as possible in negative controls, while still identifying most if not all mRNAs in donor cells. Following automatic batch analysis, each image was examined manually using the spot inspector tool to exclude obvious false positive spots. Examples of false positive spots include those at the contact sites between donor and acceptor cells (*i.e*. if cell outlining was inaccurate), spots surrounding the nucleus (as the nuclear outline was not always perfectly drawn), and spots at obvious regions of high auto-fluorescence. Thus, our analysis is generally an underestimation of the actual number of transferred mRNAs due to the abovementioned reasons. In particular, highly abundant mRNAs (*e.g.* β-actin and HER2) are underestimated in donor cells due to areas of crowding that do not allow for single molecule resolution under our imaging conditions.

Spot analysis using Airlocalize was performed as previously described (22). Briefly, high intensity pixels were detected by 3D bandpass filtering of the image. A threshold was applied so that negative control samples (*i.e.* WT MEFs alone) showed near-zero detected spots, with a minimal loss of observable spots in donor MBS MEFs.

Spot analysis using rajlabimagetools software package (available here: <u>https://bitbucket.org/arjunrajlaboratory/rajlabimagetools/wiki/Home</u>) was performed as previously described (39). Spot detection consisted of first applying a threshold across both channels that the RNA were labeled with. From there, co-localization analysis was performed to find spots that co-localized across both channels, providing an extra layer of specificity. Finally, every co-localized RNA spot corresponding to a putative RNA transfer event was examined manually and annotated to exclude any artifacts.

#### Mycoplasma infections

DAPI staining in FISH experiments was used to routinely check for mycoplasma infections. Infection was detected only once in a U2OS cell-line, and the cells were treated for 3 weeks with 25µg/ml Plasmocin (InvivoGen) until curing was observed.

### SI Figure Legends

**Figure S1.**
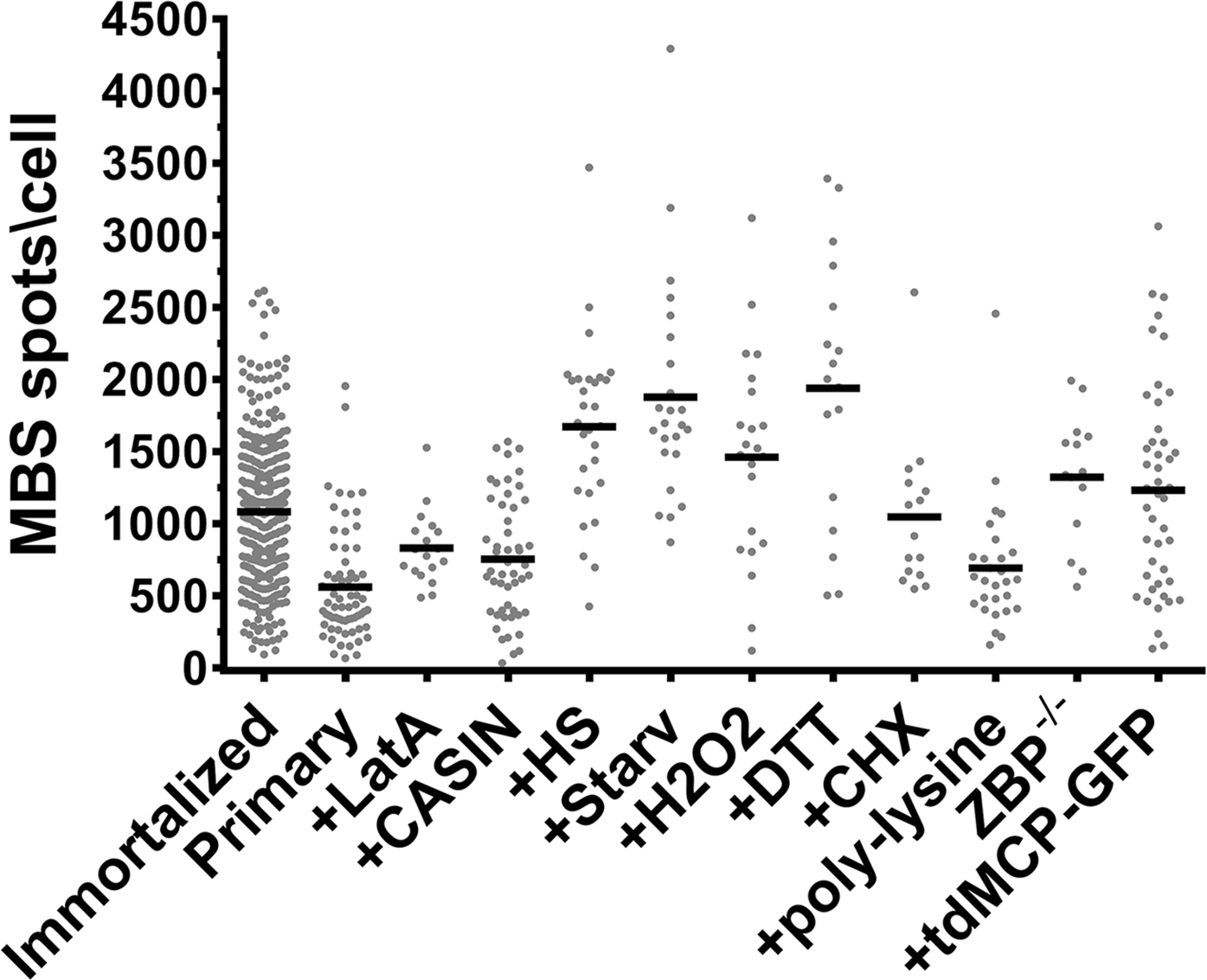
β-actin-MBS mRNA expression levels in donor MBS MEFs. Distribution of the number of β-actin-MBS mRNA (MBS) spots in immortalized MBS MEFs, primary MBS MEFs or immortalized MBS MEFs cultured under the different conditions described in the paper, as scored by smFISH using MBS probes. The bar indicates the arithmetic mean of MBS spots per donor cell per experimental condition. Conditions included: added Latrunculin A (+LatA), CASIN (+CASIN), heatshock (+HS), serum starvation (+Starv), hydrogen peroxide (+H2O2), dithiothreitol (+DTT), cycloheximide (+CHX), poly-lysine (+poly-lysine), and the absence of ZBP1 (ZBP^-/-^) or presence of tandem MCP-GFP (+tdMCP-GFP).

**Figure S2.**
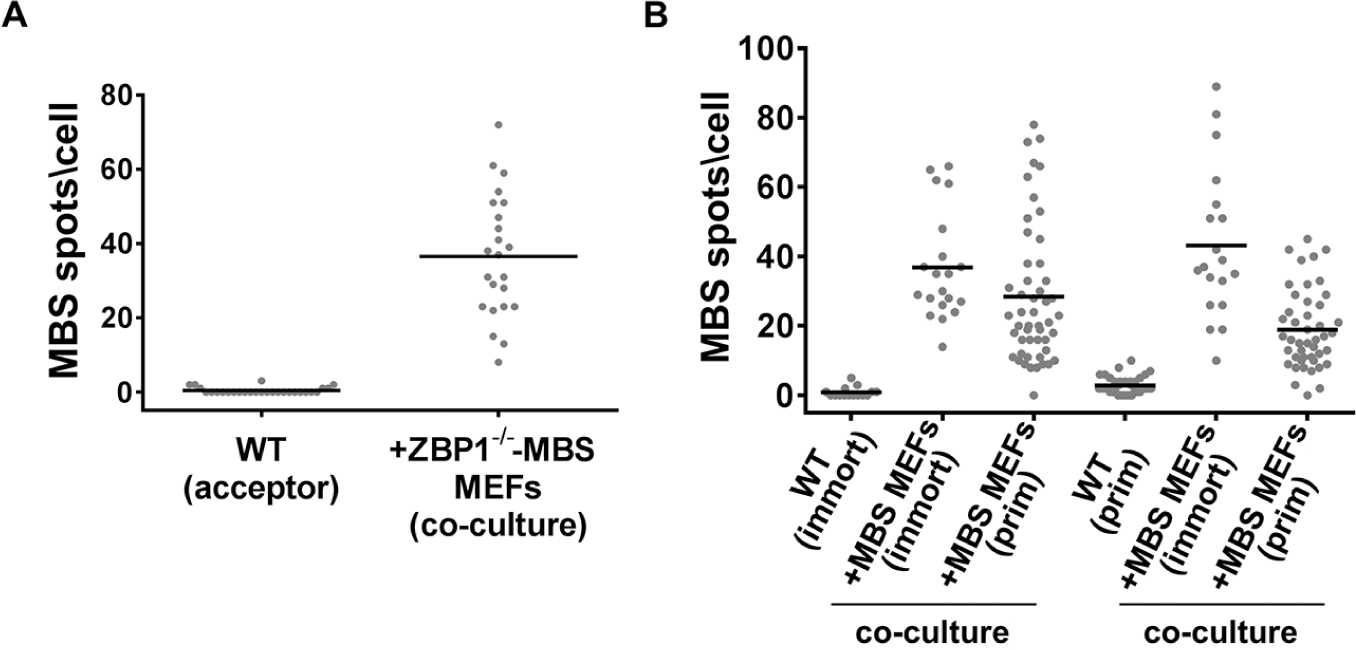
β-actin-MBS mRNA transfer in ZBP1^-/-^ and primary MEFs. **(A)** Distribution of the number of β-actin-MBS mRNA spots in acceptor WT MEFs co-cultured for 24hrs with donor ZBP1^-/-^-MBS MEFs, as scored by smFISH using MBS probes. Bar: mean of MBS spots per acceptor cell. **(B)** Distribution of the number of β-actin-MBS mRNA spots in immortalized (immort) or primary (prim) acceptor WT cells cultured alone or co-cultured for 2.5hrs with donor iMBS or pMBS MEFs, as scored above. Bar: mean number of MBS spots per acceptor cell.

**Figure S3.**
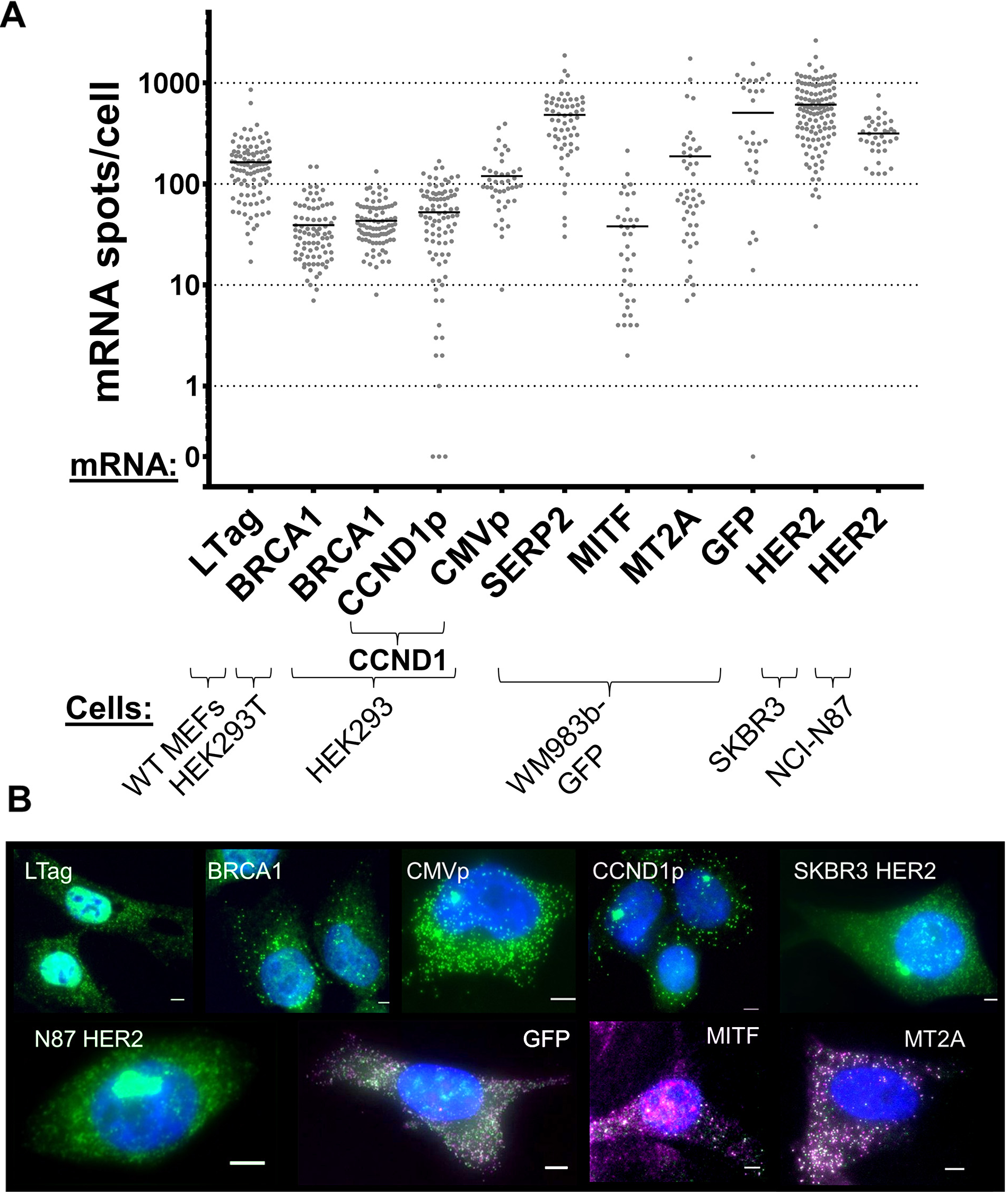
mRNA expression levels in donor cells. **(A)** Distribution of the number of mRNA spots of the different studied mRNAs in the various donor cells, as listed. Note that the Y-axis is logarithmic scale. Scoring was performed using smFISH with specific probes, listed in B. Bar: mean of mRNA spots per cell. **(B)** Representative FISH images of donor cells. Blue – DAPI labeling of the nucleus. Green – tiled FISH probes against LTag, human BRCA1, MBS (CCND1p or CMVp-CCND1-MBS), and human HER2. Green/magenta – evens/odds probe sets against GFP, human MITF and human MT2A. Scale bars: 5µm.

**Figure S4.**
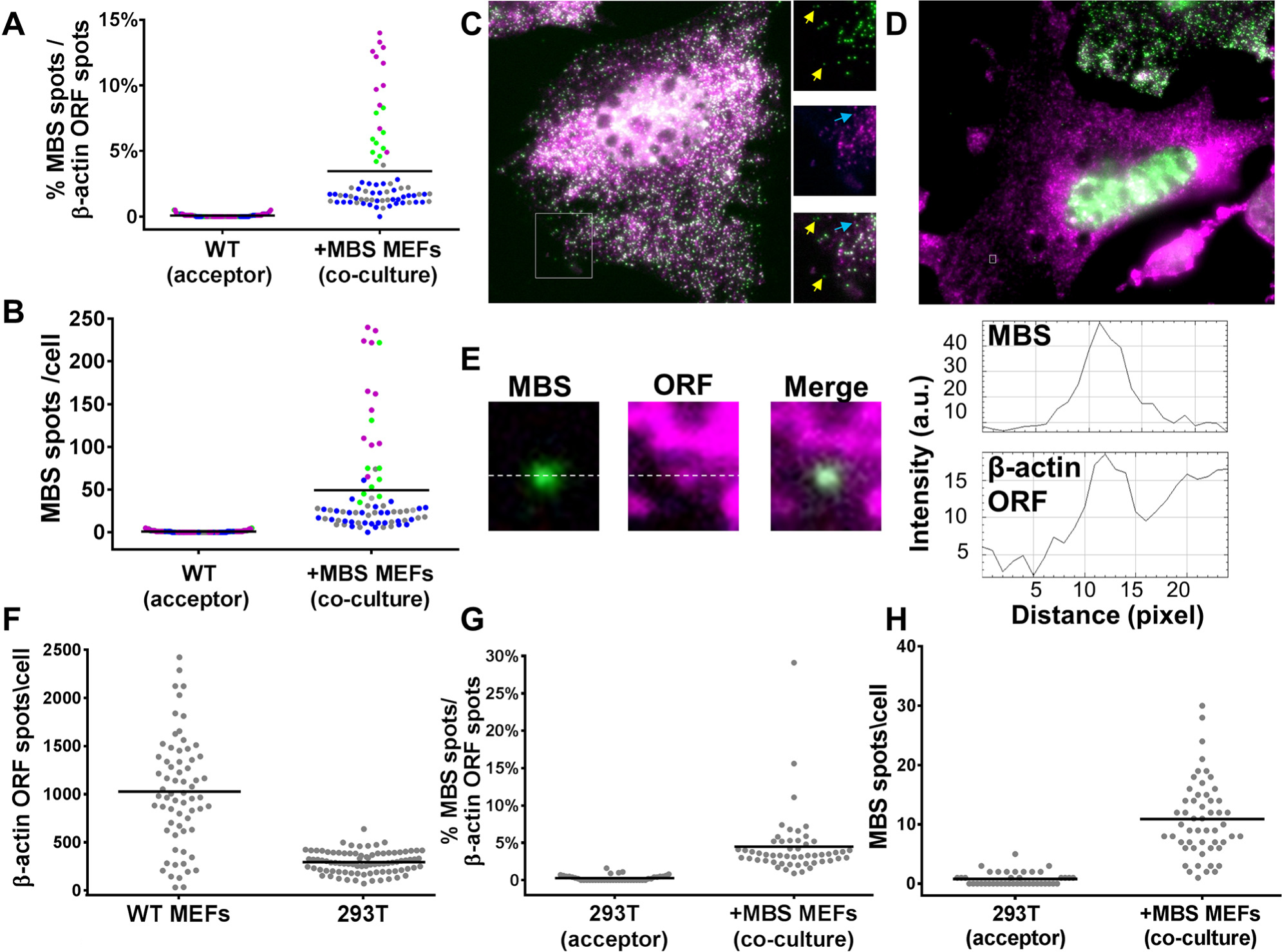
Co-FISH of β-actin-MBS mRNA transfer. **(A)** Percentage of transferred β-actin-MBS mRNA molecules out of the total β-actin mRNA population in acceptor cells. Transferred β-actin-MBS mRNA (detected with Cy3-labeled MBS probes) was scored in comparison to the total population of β - actin ORF mRNA (detected with Cy5-labeled β-actin ORF probes). Colored dots indicate results from different co-culture experiments, as follows: WT MEFs and donor MBS MEFs co-cultured for 3hrs (green), 3.5hrs (gray), and 6hrs (blue); or APAF-1^k/o^ MEFs and donor MBS MEFs co-cultured for 3hrs (magenta). Bar: mean percentage of transferred β-actin-MBS mRNA spots out of the total β-actin ORF mRNA spots in acceptor cells. **(B)** Distribution of the number of β-actin-MBS mRNA spots in acceptor MEFs following co-culture. The amount of transferred β-actin-MBS mRNA (detected with Cy3-labeled MBS probes) in the experiments shown in panel *A* was scored. Color-coding of the dots is the same as described in *A*. Bar: mean of transferred MBS mRNA spots per acceptor cell. **(C-D)** Images of co-FISH with probes against MBS (green) and β-actin ORF (magenta). A co-labeled donor MBS MEF is shown in *C*. Note the numerous MBS spots (green) and ORF spots (magenta). Panels to the right show zoom-in images of the rectangle found in the main panel (left). Yellow arrows indicate MBS spots that do not co-localize with ORF probes. Blue arrows indicate ORF spots that do not co-localize with the MBS spots. An acceptor WT MEF in co-culture is shown in the center of *D*. Note the low amounts of MBS spots (green) in comparison to ORF spots (magenta). **(E)** An example of a transferred β-actin-MBS mRNA spot (MBS; green) from *D* co-localized with a β-actin ORF spot (ORF; magenta) from the cell in panel *D*. The adjacent graphs show pixel intensity across the white dotted median. **(F)** Distribution of the number of β-actin mRNA molecules in WT MEFs and HEK293T cells by using the β-actin ORF probe set. Bar: mean number of spots per cell **(G)** Percentage of transferred β-actin-MBS mRNA molecules out of the total β - actin mRNA in acceptor HEK293T cells. Scoring was performed as in *A.* **(H)** Distribution of the number of transferred β-actin-MBS mRNA spots in acceptor HEK293T cells following co-culture with MBS MEFs. The amount of transferred β-actin-MBS mRNA (detected with Cy3-labeled MBS probes) in the experiment shown in panel *G* was scored. Bar: mean number of transferred mRNA spots per acceptor cell.

**Figure S5.**
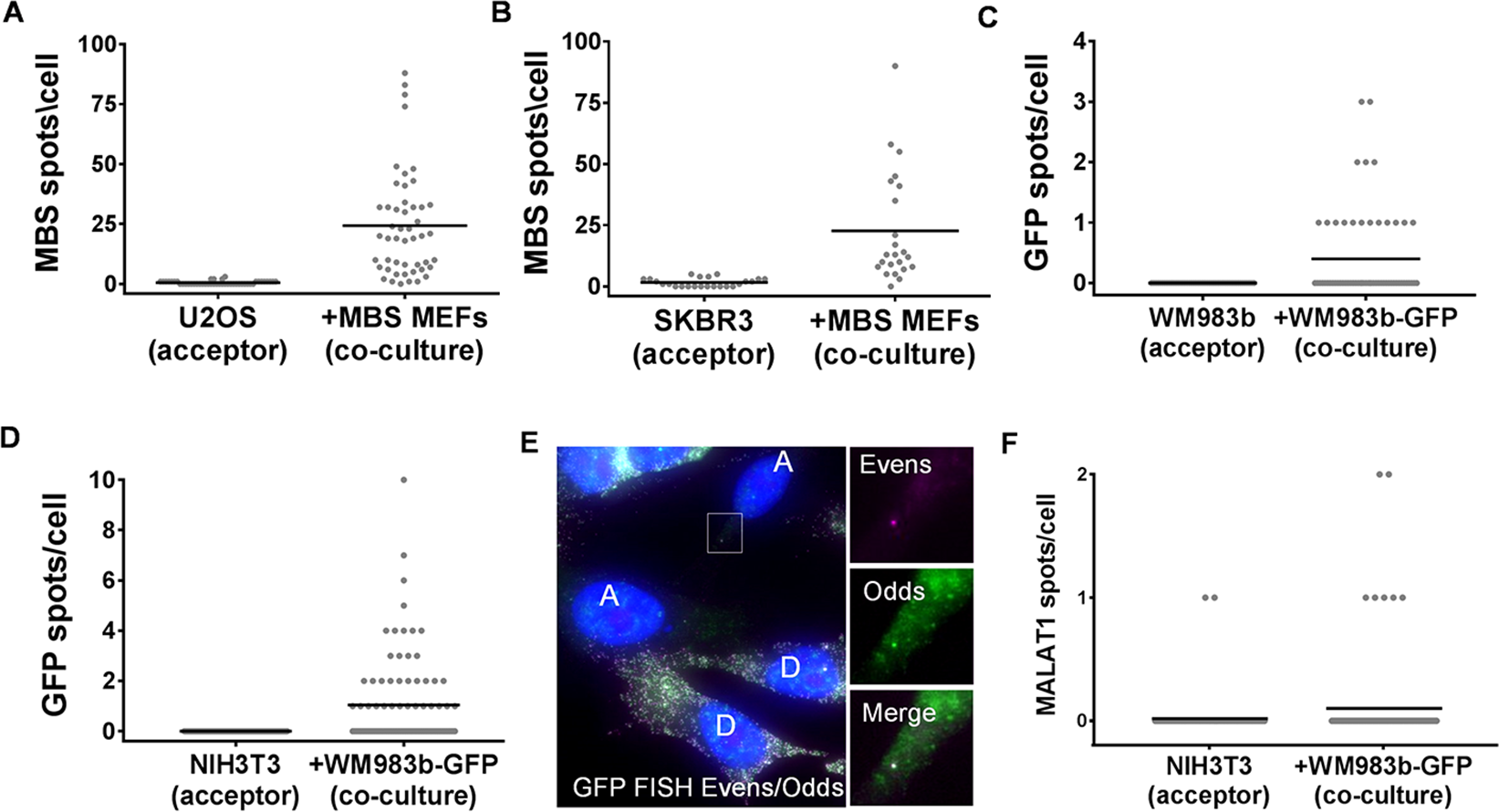
Transfer of mRNA in a human-murine co-culture. **(A-B)** Distribution of the number of β - actin-MBS mRNA spots in U2OS cells (A) or SKBR3 cells (B) that were co-cultured with donor MBS MEFs for 24hrs. Bar: mean of transferred β-actin-MBS mRNA spots per acceptor cell. **(C)** Distribution of the number of GFP mRNA spots in human melanoma cells WM983b that were co-cultured with donor WM983b-GFP cells for 48hr. Bar: mean of transferred GFP mRNA spots per acceptor cell. Only dual-color spots were considered as legitimate mRNA spots. **(D)** Distribution of the number of GFP mRNA spots in NIH3T3 cells that were co-cultured with donor WM983b-GFP cells for 48hr. Bar: mean of transferred GFP mRNA spots per acceptor cell. Only dual-color spots were considered as legitimate mRNA spots. **(E)** Image of co-FISH with odds/evens probes against GFP. D indicates donor WM983b-GFP cells, A indicates WM983b acceptor cells. Magenta label - Alexa594 (evens), green label - Cy3 (odds). Zoom-in photos of the boxed area in the main panel (left) are shown (right). **(F)** Distribution of the number of MALAT1 RNA spots in NIH3T3 cells that were co-cultured with donor WM983b-GFP for 48hrs. Bar: mean of transferred MALAT1 RNA spots per acceptor cell.

**Figure S6.**
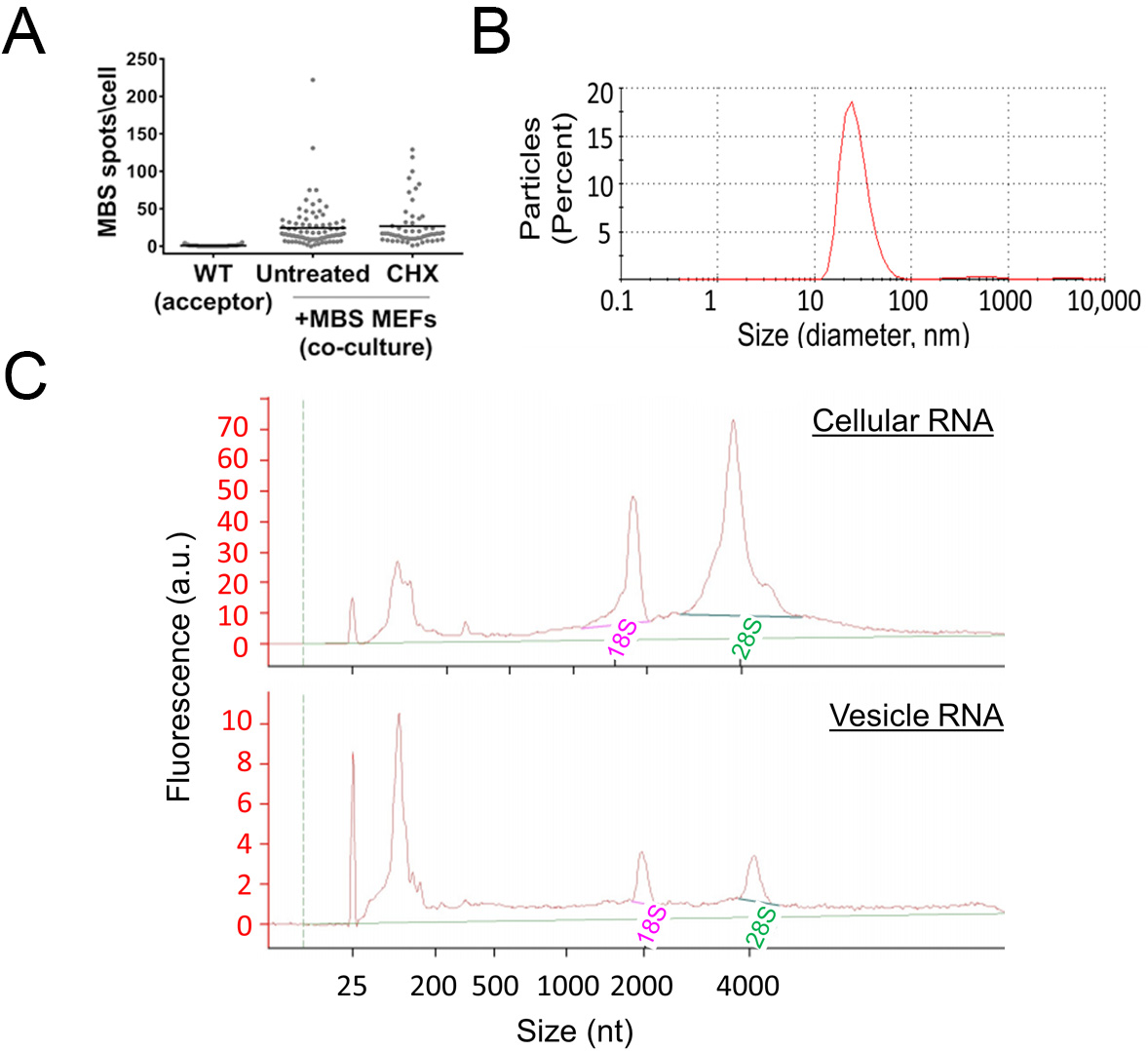
Translation inhibition does not inhibit β-actin-MBS mRNA transfer; isolated exosomes contain primarily small RNAs. Acceptor WT MEFs were first co-cultured (0.5hrs) with donor MBS MEFs, followed by additional co-culture for 2.5hrs in the absence (Untreated) or presence of cycloheximide (CHX; 100µg/ml). FISH analysis of β-actin-MBS mRNA was performed using Cy3-labeled MBS probes. Bar: mean of transferred MBS mRNA spots per acceptor cell. **(B)** Exosomes size was measured as described in the Methods section. **(C)** RNA isolated from whole cell lysate (Cellular RNA) or exosomes (Vesicles RNA) was analyzed using a Bioanalyzer. 18S and 28S indicate peaks of ribosomal RNAs.

**Figure S7.**
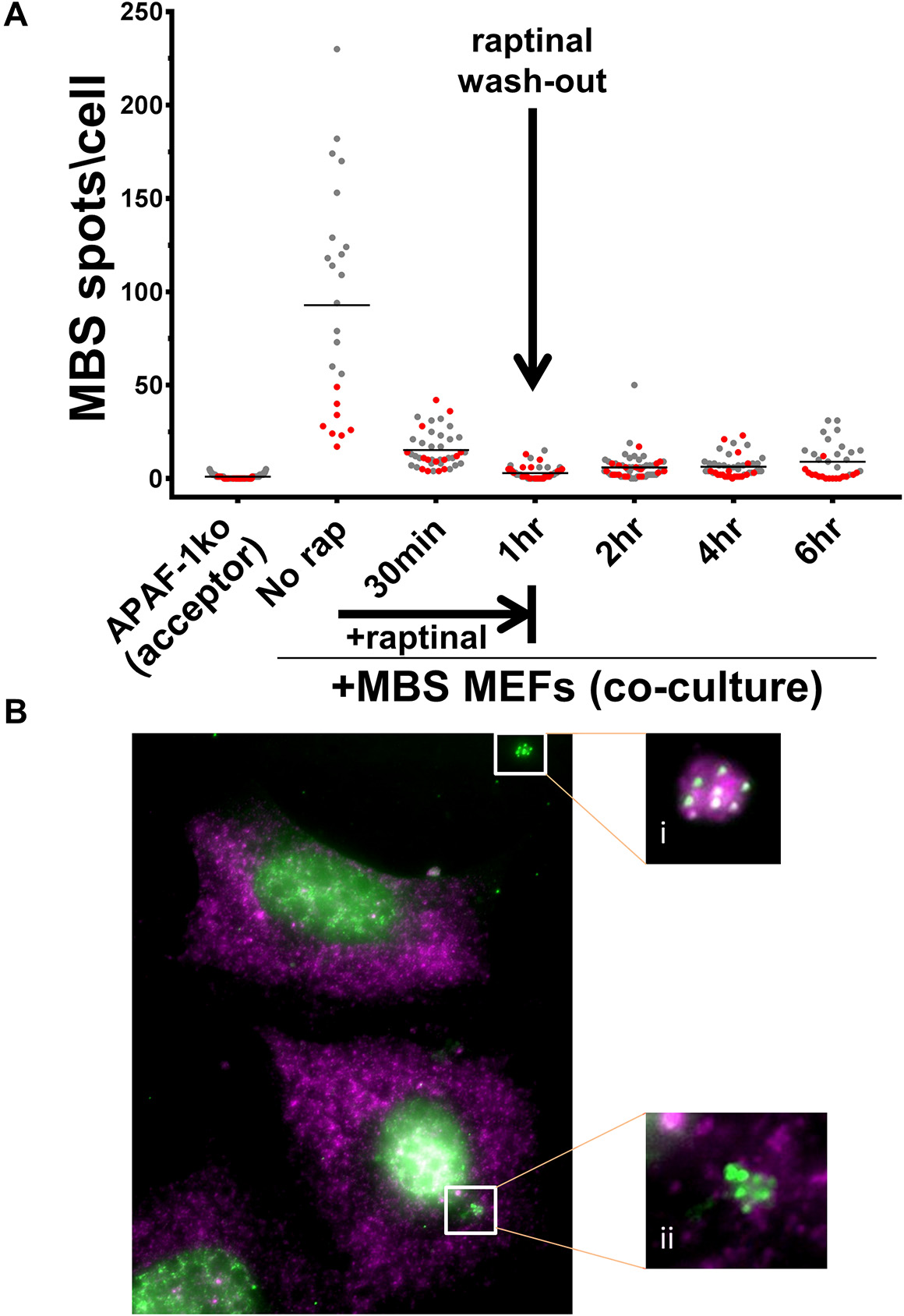
Cell death does not contribute significantly to mRNA transfer. **(A)** APAF-1^k/o^ MEFs were co-cultured with donor MBS MEFs. Cells were either co-plated and cultured together for 12hrs (red spots) or MBS MEFs were plated on top of pre-seeded (overnight) APAF-1^k/o^ MEFs and co-cultured for 3hrs (gray spots). Following co-culture, cells were treated with raptinal for 1hr. smFISH for β-actin-MBS mRNA using Cy3-labeled probes was performed at 0.5,1,2,4, and 6hrs after the addition of raptinal. Bar: mean of transferred β-actin-MBS mRNA spots per acceptor cell. **(B)** Image of co-FISH with probes against MBS (Cy3 label, green) and β-actin ORF (Cy5 label, magenta) in acceptor APAF-1^k/o^ MEFs at 4hrs (*i.e.* 3hrs after raptinal wash-out). White boxes show extracellular (i) and intracellular (ii) apoptotic bodies containing MBS spots.

**Figure S8.**
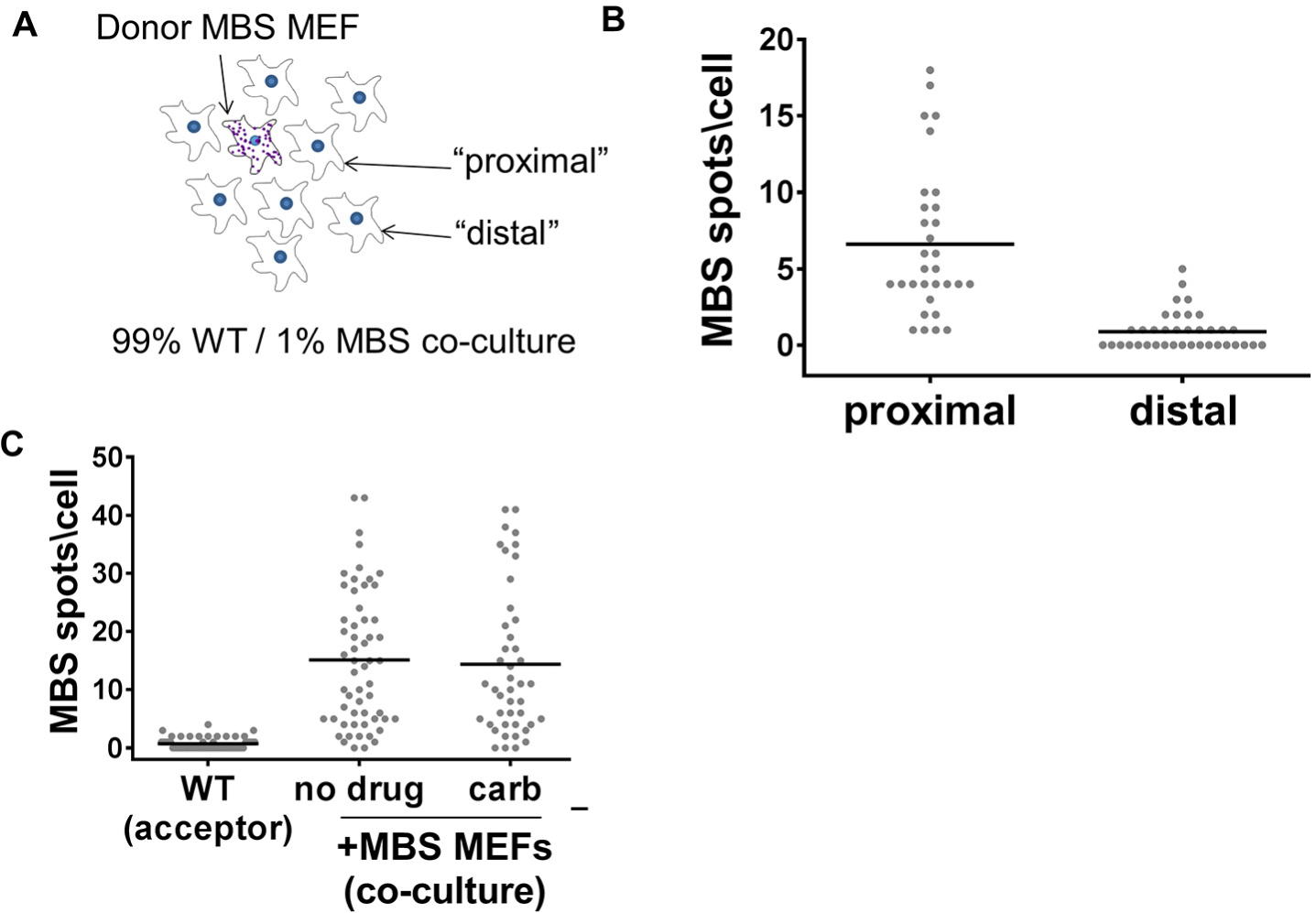
mRNA transfer requires proximity between cells, but does not occur through Gap-junctions. **(A)** Illustration of the 99% WT MEF::1% MBS MEF co-culture system. Acceptor cells immediately surrounding the donor MBS MEF (*i.e.* first cell layer) are designated as “proximal” cells, while acceptors located beyond the first layer are labeled as being “distal”. **(B)** Distribution of the number of β-actin-MBS mRNA spots in 99% WT MEF::1%MBS MEF co-cultures. Donor MBS MEFs and acceptor MEFs were plated simultaneously at a 1:99 ratio and were co-cultured for 24hrs, prior to scoring MBS spots in acceptor cells by smFISH using MBS probes. The number of transferred MBS spots was determined in proximal vs. distal acceptor cells. Bar: mean of transferred MBS mRNA spots per proximal or distal acceptor cell. **(C)** Use of a Gap junction inhibitor does not block mRNA transfer. Acceptor WT MEFs and donor MBS MEFs were co-cultured for 2hrs in the absence (no drug) or presence of Carbenoxolone (carb), followed by smFISH analysis of β-actin-MBS mRNA. Bar: mean of transferred MBS mRNA spots per acceptor cell.

**Figure S9:**
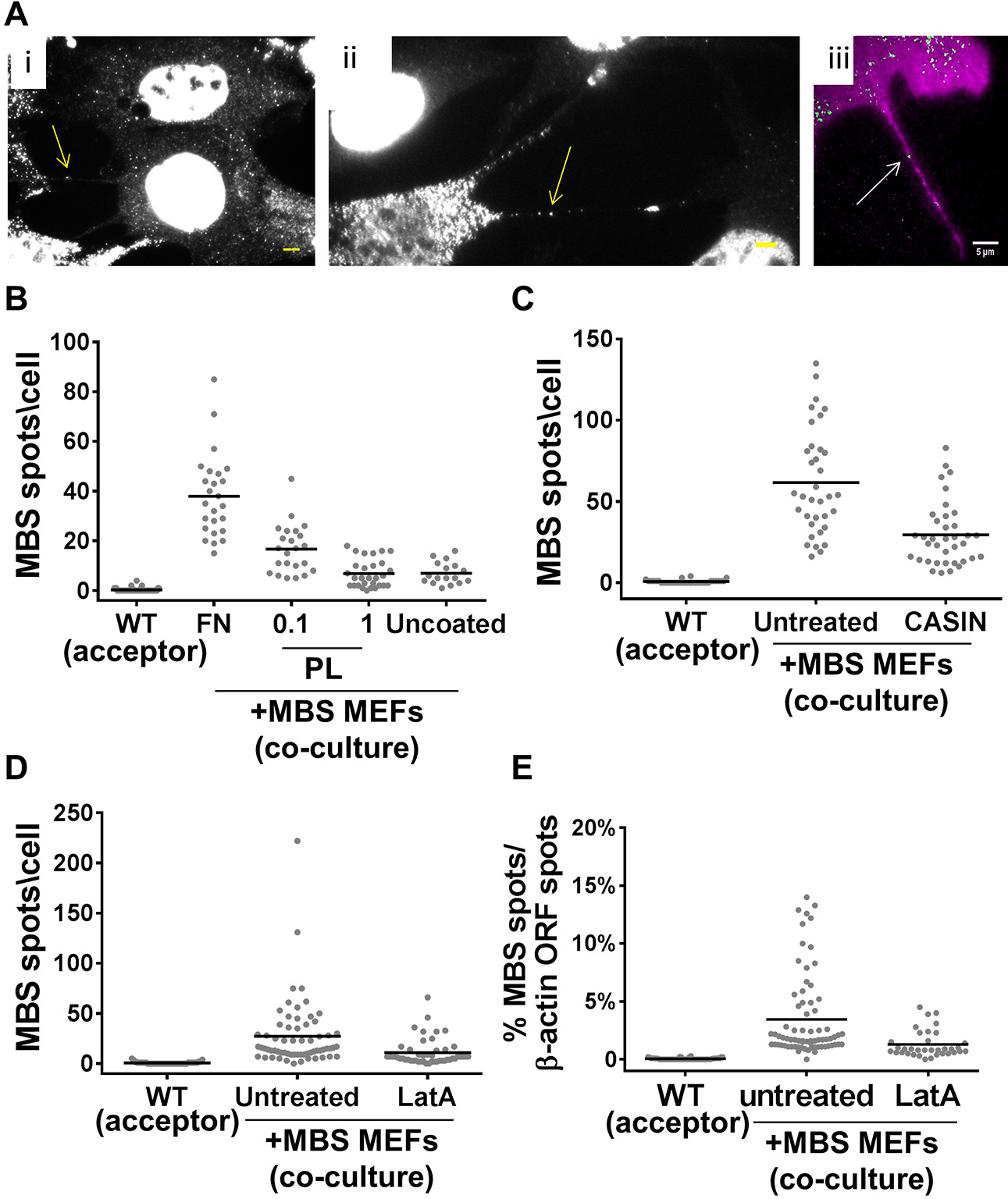
Treatment with mNT inhibitors reduces mRNA transfer. **(A)** β-actin-MBS mRNA is present in mNTs. **(i)** FISH image of a β-actin-MBS mRNA spot present in a mNT formed between two cells. Scale bar: 5µm. **(ii)** FISH image of β-actin-MBS mRNAs in a mNT formed by a MBS MEF. Scale bar: 5µm. **(iii)** Live image of a β-actin-MBS mRNA in a mNT produced by a MBS MEF. Arrow points to a tdMCP-GFP-labeled mRNA (green). Both cell types were labeled with membrane-targeted TagRFP-T-ps (magenta). TagRFP-T-ps- and tdMCP-GFP-expressing MBS MEFs were plated, co-cultured with TagRFP-T-ps-expressing WT MEFs (not shown) for 2-3hrs, and then imaged using a 150x objective, as detailed in the Materials and Methods (see Live Imaging section). Cells were imaged at 8 *z*-sections of 0.6µm each. Exposure time at each wavelength was 100ms. The 488nm laser was set at 20% power; 561nm laser at 10% power. Scale bar: 5µm. **(B)** Fibronectin promotes mRNA transfer. WT MEFs were cultured on uncoated glass or on either fibronectin (FN)- or poly-lysine (PL; 0.1mg/ml or 1mg/ml)-coated glass, prior to co-culture with donor MBS MEFs for 2.5hrs, and then analyzed for β-actin-MBS transfer by smFISH with MBS probes. Bar (in panels *B-D*): mean of transferred MBS mRNA spots per acceptor cell. **(C)** CASIN inhibits the transfer of β-actin-MBS mRNA. Acceptor WT MEFs were first co-cultured (0.5hrs) with donor MBS MEFs alone, followed by the addition of CASIN (1µM) and continued co-culture for 2.5hrs, prior to smFISH analysis using MBS probes. **(D)** Latrunculin A inhibits the transfer of β-actin-MBS mRNA. Acceptor WT MEFs were first co-cultured (0.5hrs) with donor MBS MEFs alone, followed by the addition of Latrunculin A (LatA; 200nM) and continued co-culture for 2.5hrs, prior to smFISH analysis using MBS and β-actin ORF probes. Scoring of β-actin-MBS spots is shown. **(E)** Percentage of transferred β-actin-MBS mRNA molecules out of total β-actin mRNA in LatA-treated acceptor cells. Untreated and LatA-treated acceptor WT MEFs from panel *D* were scored for the percentage of β-actin-MBS mRNAs (measured using Cy3-labeled MBS probes) out of the total β-actin mRNA pool (measured using Cy5-labeled ORF probes) using smFISH. Bar: mean percentage of transferred MBS mRNA spots out of total β-actin pool per acceptor cell.

**Figure S10.**
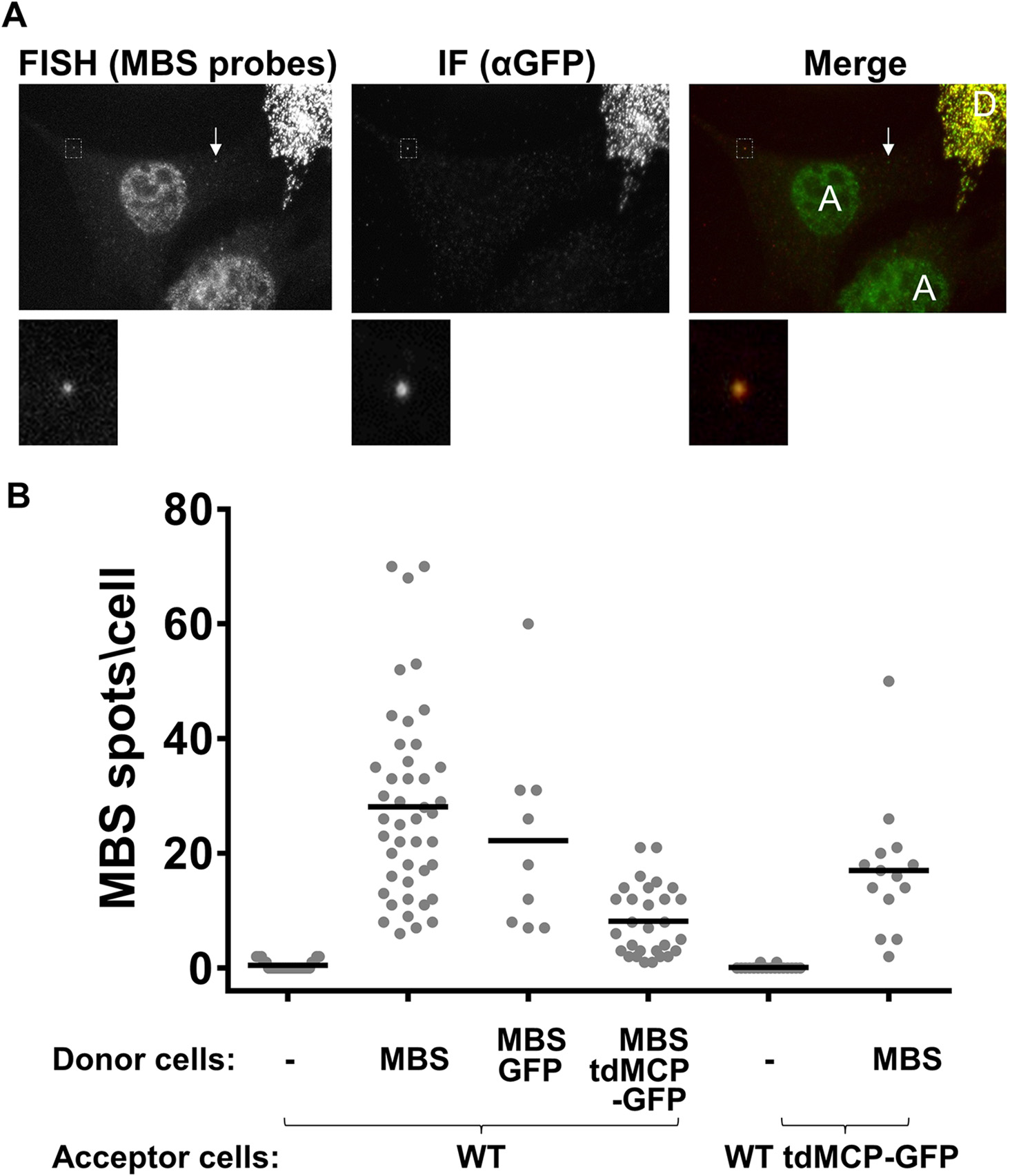
The MS2 coat protein inhibits MBS-labeled mRNA transfer. **(A)** FISH-IF images of WT and MBS MEFs expressing tdMCP-GFP. Acceptor WT MEFs (labeled as *A*) were co-cultured with donor MBS MEFs expressing tdMCP-GFP (labeled as *D*). Cells were co-cultured for 2.5hrs prior to FISH-IF using MBS probes to detect β-actin-MBS mRNA (green in *merge*) and chicken anti-GFP antibodies to detect tdMCP-GFP (red in *merge*). Top row images show co-localization of most FISH spots with immunofluorescence labeling in the donor cell and a single transferred mRNA in the acceptor cell. Arrow indicates a transferred mRNA without tdMCP-GFP labeling. Bottom row images show a higher magnification of the boxed FISH- and immunofluorescence-labeled spot in the top panels. **(B)** β-actin-MBS mRNA transfer is inhibited by donor cells expressing tdMCP-GFP. Acceptor WT MEFs (WT) or acceptor WT MEFs expressing tdMCP-GFP (WT tdMCP-GFP) were co-cultured for 2.5hrs with donor MBS MEFs alone (MBS) or donor MBS MEFs expressing either GFP (MBS GFP) or tdMCP-GFP (MBS tdMCP-GFP). mRNA transfer was scored by smFISH MBS using MBS probes. Bar: mean number of transferred MBS mRNA spots per acceptor cell.

## SI Table Legend

**Table S1.** The spreadsheet presents the following data for each graph/column: *N* – number of cells; *average* – calculated mean of the data in each column; *SEM* – standard error of the mean; *p*-value – as determined by unpaired t-tests between the indicated sets of results.

### SI Movie Legends

**Movie S1.**
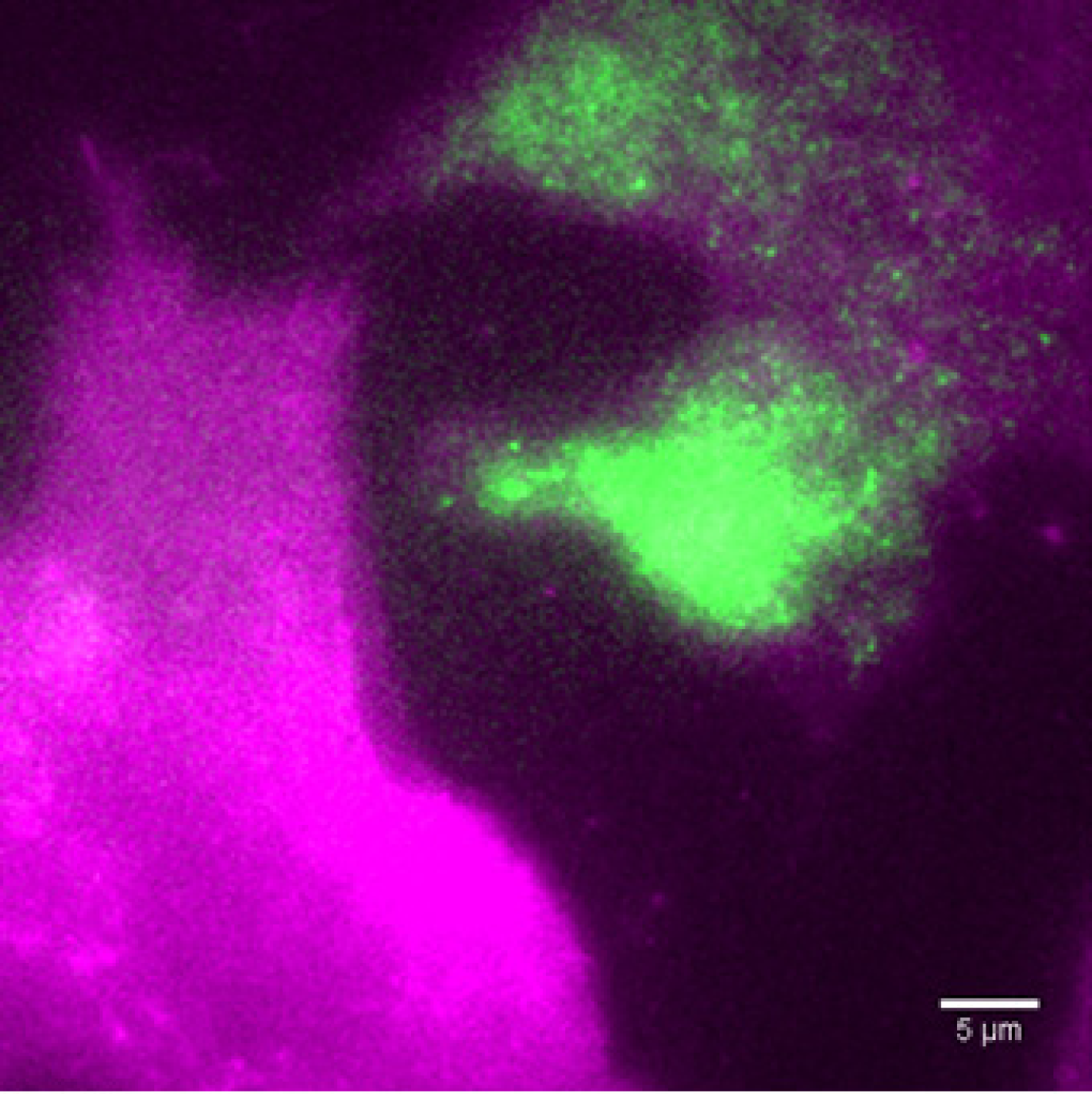
Transfer of β-actin-MBS mRNA between donor and acceptor MEFs. Acceptor WT MEFs and donor β-actin-MBS MEFs expressing tdMCP-GFP were co-cultured for 2.5hrs and imaged with a 150x objective, as detailed in the Materials and Methods section. Both cell types express TagRFP-T-ps. Cells were imaged at 7sec intervals, 8 *z*-sections of 0.6µm each, for 14min. Exposure time for each wavelength was 100ms (488nm laser at 20% power, 561nm laser at 20% power). Green – GFP, Magenta – TagRFP-T. Movie shows maximum projection of 61 frames at 5fps (total elapsed time = ~7min, starting from frame #30) of the upper two *z*-sections. Scale bar = 5µm. Note the linear progress of the β-actin-MBS mRNA spot between donor and acceptor MEFs.

**Movie S2.**
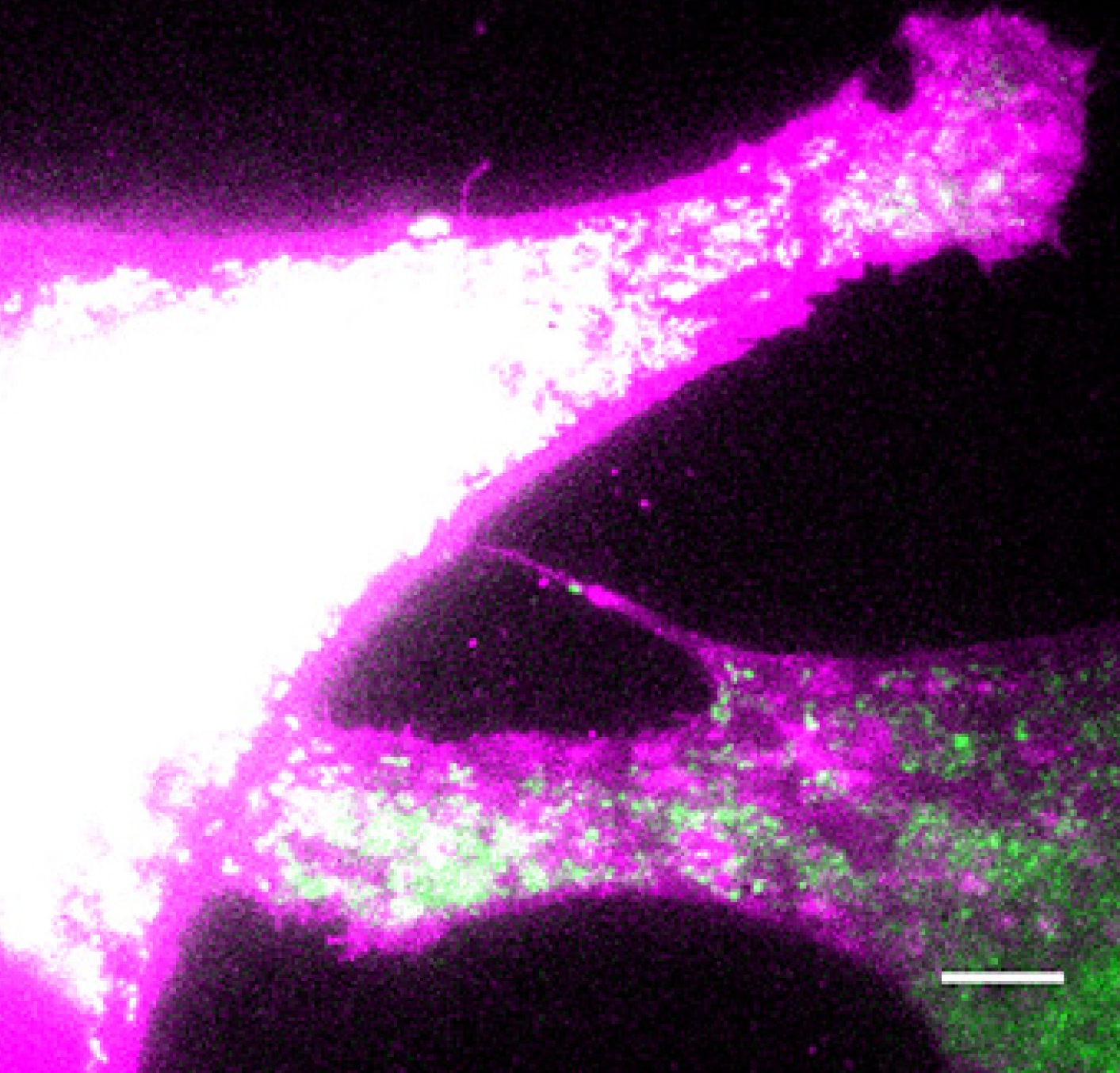
Transfer of β-actin-MBS mRNA between donor MEFs. Donor β-actin-MBS MEFs expressing tdMCP-GFP and TagRFP-T-ps were co-cultured (with WT MEFs; not shown) for 2.5hrs and imaged with a 150x objective, as detailed in the Materials and Methods section. Cells were imaged at 10sec intervals, 11 *z*-sections of 0.6µm each, for 20min. Exposure time for each wavelength was 300ms (488nm laser at 50% power, 561nm laser at 20% power). Green – GFP, Magenta – TagRFP-T. Movie shows maximum projection of each of the 121 frames at 5fps. Scale bar = 5µm. Note that the donor MBS cell on the right makes two polarized membrane extensions. The first (at the center of the frame) is a mNT that extends out to the acceptor MBS cell on the left (and contains a β-actin-MBS mRNA spot), while the second (at the bottom-left of the frame) is a lamellipodial-like extension. *z*-stack analysis reveals that the second extension likely makes contact with the left cell, but does not result in a cytoplasmic connection between the two cells.

**Movie S3.**
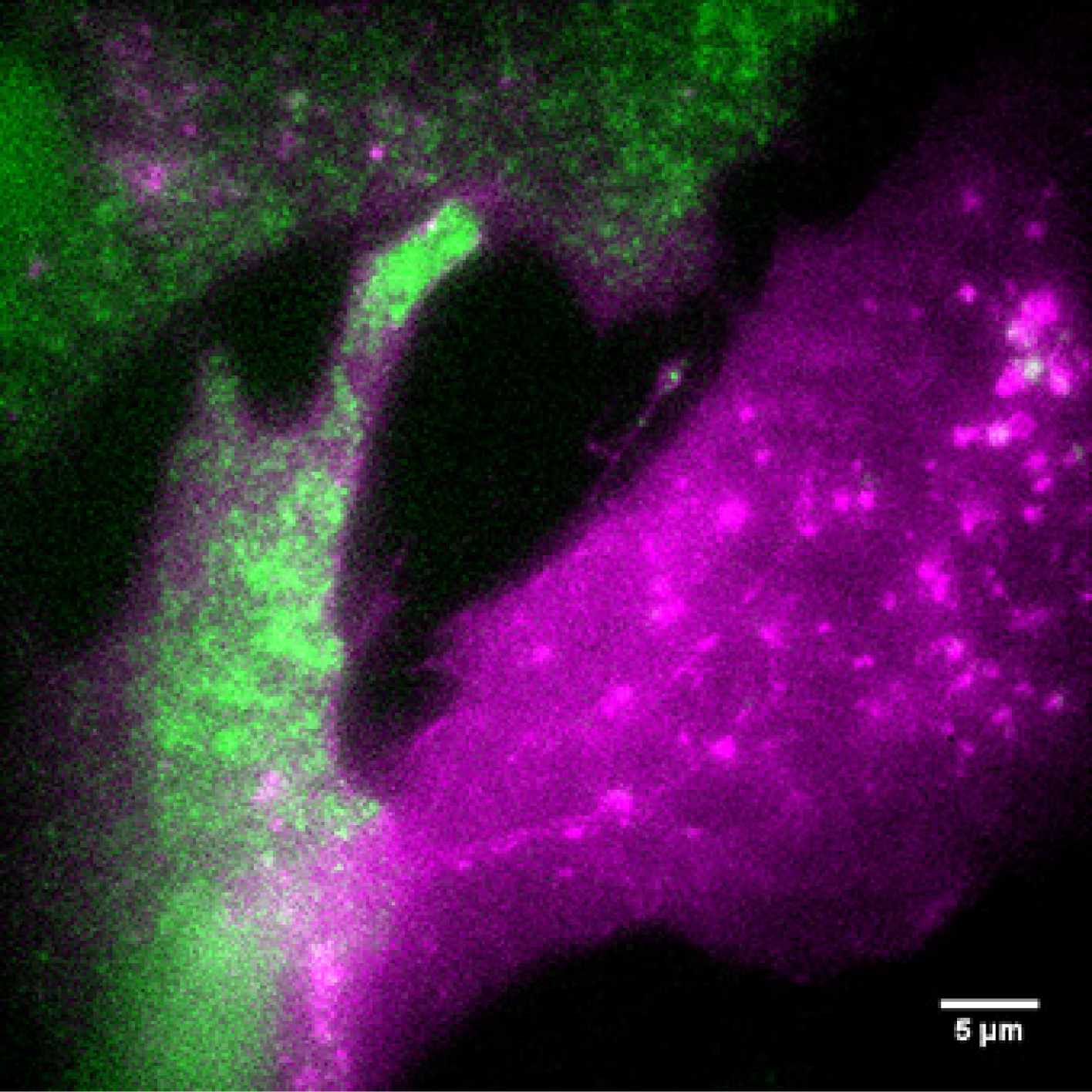
Transfer of β-actin-MBS mRNA between donor and acceptor MEFs. Acceptor WT MEFs and donor β-actin-MBS MEFs expressing tdMCP-GFP (green) were co-cultured for 2.5hrs, and then imaged with 60x objective, as detailed in the Materials and Methods section. Both cell types express TagRFP-T-ps (magenta). Cells were imaged at 3sec intervals, 8 *z*-sections of 0.6µm each, for 3min. Exposure time for each wavelength was 100ms (488nm laser at 20% power, 561nm laser at 10% power). Movie shows maximum projection of each of 121 frames at 5fps. Scale bar, 5µm. Note the mNT with GFP-labeled β - actin-MBS spots in the center of the movie. Also note the physical contact, but lack of GFP transfer, between the right WT MEF and the left MBS MEF.

**Movie S4.**
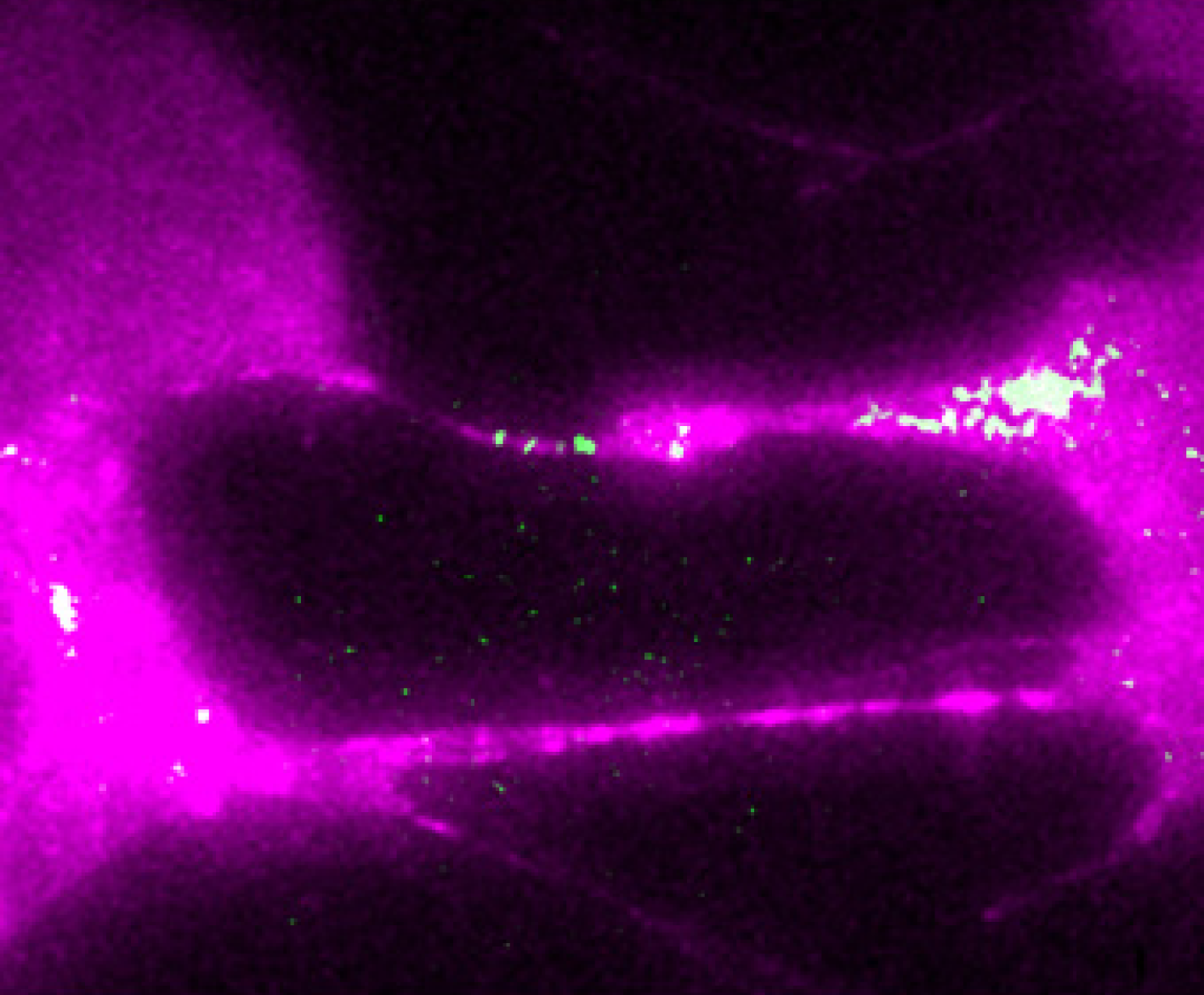
Transfer of β-actin-MBS mRNA between donor and acceptor MEFs. Acceptor WT MEFs and donor β-actin-MBS MEFs expressing tdMCP-GFP (green) were co-cultured for 24hrs and then imaged with a 150x objective, as detailed in the Materials and Methods section. Both cell types express TagRFP-T-ps (magenta). For this movie, cells were imaged at 10sec intervals, 11 *z*-sections of 0.6µm each, for 45min. Exposure time and laser power for each wavelength was 100ms at 50% power for the 488nm laser and 200ms at 30% power for the 561nm laser. Movie shows a maximum projection of each of 122 frames (elapsed time = 20min 10sec, starting from frame #1) at 5fps. Scale bar, 5µm. Note the presence of GFP-labeled β-actin-MBS spots in the mNT and possible retrograde transport back to the original donor cell (right).

**Movie S5.**
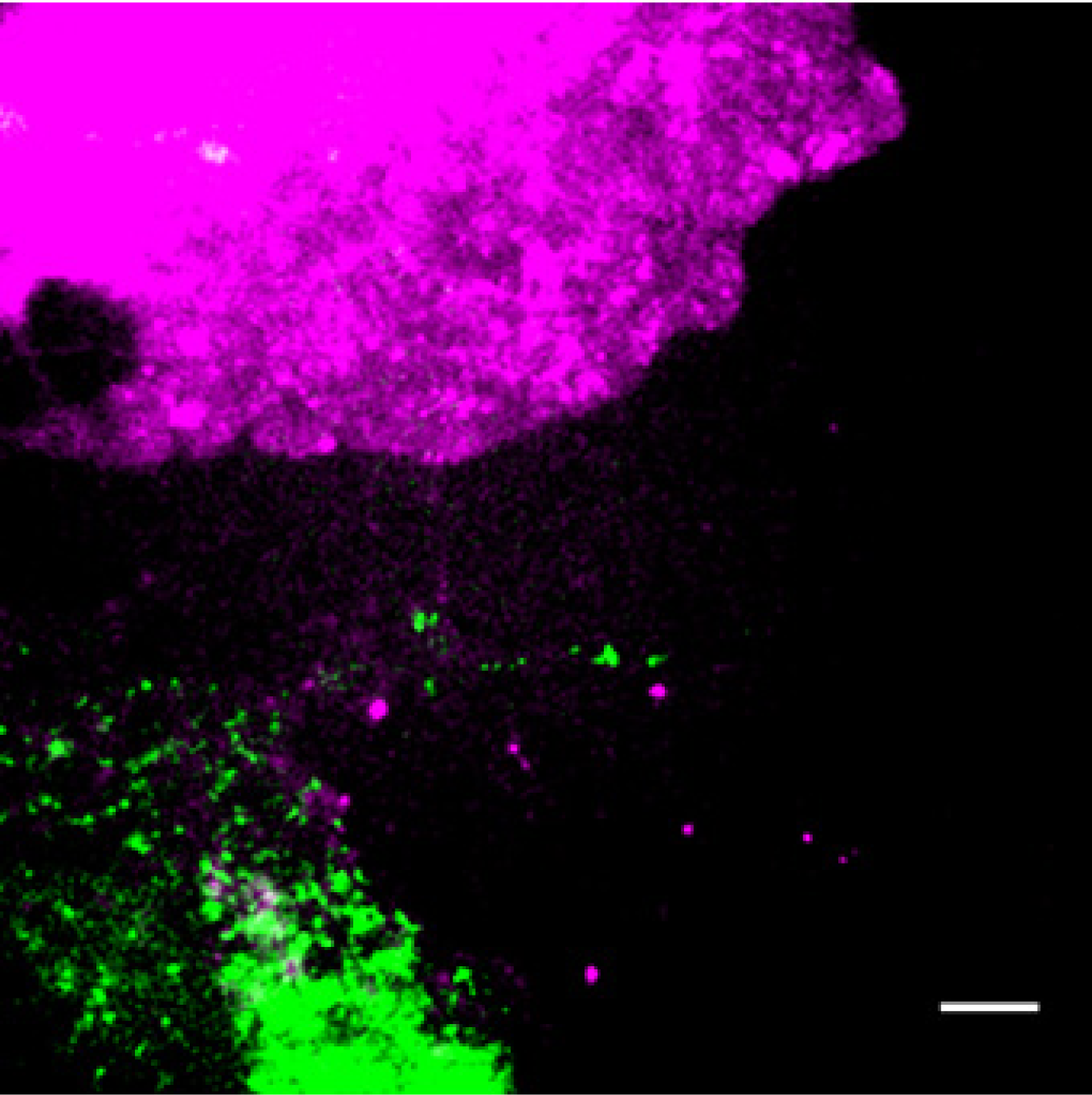
Transfer of β-actin-MBS mRNA between donor and acceptor MEFs. Acceptor WT MEFs and donor β-actin-MBS MEFs expressing tdMCP-GFP (green) were co-cultured for 2.5hrs and then imaged with a 150x objective, as detailed in the Materials and Methods section. Both cell types express TagRFP-T-ps (magenta). For this movie, cells were imaged at 10sec intervals, 11 *z*-sections of 0.6µm each, for 10min. Exposure time and laser power for each wavelength was 600ms at 55% power for the 488nm laser, 400ms at 35% power for 561nm laser. Movie shows maximum projection of each of the 7 frames (frames 48-54) at 1fps. Scale bar, 5µm. Note the apparent transfer of a GFP-labeled β-actin-MBS spot from the lower cell via the mNT and its appearance in the cytoplasm of the upper acceptor cell.

**Movie S6.**
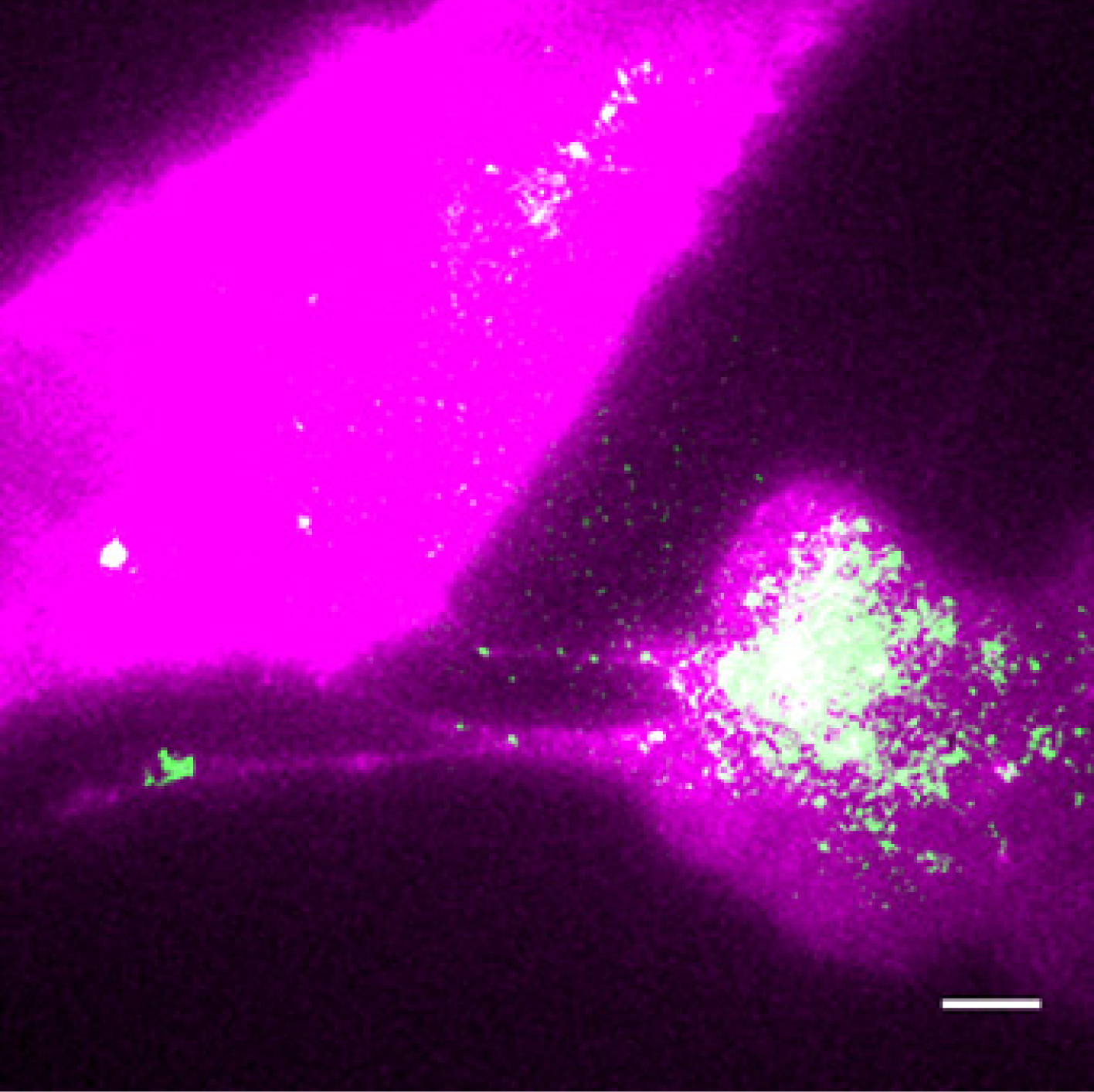
Transfer of β-actin-MBS mRNA between heterologous donor and acceptor cell types. Acceptor U2OS cells and donor β-actin-MBS MEFs expressing tdMCP-GFP (green) were co-cultured for 2.5hrs and then imaged with a 150x objective, as detailed in the Materials and Methods section. Both cell types express TagRFP-T-ps (magenta). For this movie, cells were imaged at 10sec intervals, 11 *z*-sections of 0.6µm each, for 30min. Exposure time and laser power for each wavelength was 100ms at 50% power for the 488nm laser and 200ms at 30% power for the 561nm laser. Movie shows a maximum projection of each of 121 frames (20min, starting from frame #1) at 5fps. Scale bar, 5µm. Note the presence of GFP-labeled β-actin-MBS spots in the mNT.

## References

[1] Kolodny GM (1971) Evidence for transfer of macromolecular RNA between mammalian cells in culture. Exp Cell Res 65(2):313–324.

[2] Kolodny GM (1972) Cell to cell transfer of RNA into transformed cells. J Cell Physiol 79(1):147–150.

[3] Kolodny GM, Culp LA, & Rosenthal LJ (1972) Secretion of RNA by normal and transformed cells. Exp Cell Res 73(1):65–72.

[4] Valadi H, et al. (2007) Exosome-mediated transfer of mRNAs and microRNAs is a novel mechanism of genetic exchange between cells. Nat Cell Biol 9(6):654–659.

[5] Arroyo JD, et al. (2011) Argonaute2 complexes carry a population of circulating microRNAs independent of vesicles in human plasma. Proc Natl Acad Sci USA 108(12):5003–5008.

[6] Eldh M, et al. (2010) Exosomes communicate protective messages during oxidative stress; possible role of exosomal shuttle RNA. PloS One 5(12):e15353.

[7] Xiao D, et al. (2012) Identifying mRNA, microRNA and protein profiles of melanoma exosomes. PloS One 7(10):e46874.

[8] Eirin A, et al. (2014) MicroRNA and mRNA cargo of extracellular vesicles from porcine adipose tissue-derived mesenchymal stem cells. Gene 551(1):55–64.

[9] Jiang H, Li Z, Li X, & Xia J (2015) Intercellular transfer of messenger RNAs in multiorgan tumorigenesis by tumor cell-derived exosomes. Mol Med Rep 11(6):4657–4663.

[10] Lee C, et al. (2012) Exosomes mediate the cytoprotective action of mesenchymal stromal cells on hypoxia-induced pulmonary hypertension. Circulation 126(22):2601–2611.

[11] Lopez-Verrilli MA, Picou F, & Court FA (2013) Schwann cell-derived exosomes enhance axonal regeneration in the peripheral nervous system. Glia 61(11):1795–1806.

[12] Tomasoni S, et al. (2012) Transfer of Growth Factor Receptor mRNA Via Exosomes Unravels the Regenerative Effect of Mesenchymal Stem Cells. Stem Cells Dev 22(5):772–780.

[13] Kanada M, et al. (2015) Differential fates of biomolecules delivered to target cells via extracellular vesicles. Proc Natl Acad Sci USA 112(12):E1433–1442.

[14] Tian T, et al. (2013) Dynamics of exosome internalization and trafficking. J Cell Physiol 228(7):1487–1495.

[15] Heusermann W, et al. (2016) Exosomes surf on filopodia to enter cells at endocytic hot spots, traffic within endosomes, and are targeted to the ER. J Cell Biol 213(2):173–184.

[16] Hung ME & Leonard JN (2016) A platform for actively loading cargo RNA to elucidate limiting steps in EV-mediated delivery. J Extracell Vesicles 5:31027.

[17] Batagov AO & Kurochkin IV (2013) Exosomes secreted by human cells transport largely mRNA fragments that are enriched in the 3'-untranslated regions. Biol direct 8:12.

[18] Chevillet JR, et al. (2014) Quantitative and stoichiometric analysis of the microRNA content of exosomes. Proc Natl Acad Sci USA 111(41):14888–14893.

[19] Stevanato L, Thanabalasundaram L, Vysokov N, & Sinden JD (2016) Investigation of Content, Stoichiometry and Transfer of miRNA from Human Neural Stem Cell Line Derived Exosomes. PloS One 11(1):e0146353.

[20] Raj A, van den Bogaard P, Rifkin SA, van Oudenaarden A, & Tyagi S (2008) Imaging individual mRNA molecules using multiple singly labeled probes. Nat Methods 5(10):877–879.

[21] Femino AM, Fay FS, Fogarty K, & Singer RH (1998) Visualization of single RNA transcripts in situ. Science 280(5363):585–590.

[22] Lionnet T, et al. (2011) A transgenic mouse for in vivo detection of endogenous labeled mRNA. Nat Methods 8(2):165–170.

[23] Abounit S & Zurzolo C (2012) Wiring through tunneling nanotubes-from electrical signals to organelle transfer. J Cell Sci 125(Pt 5):1089–1098.

[24] Biran A, et al. (2015) Senescent cells communicate via intercellular protein transfer. Genes Dev 29(8):791–802.

[25] Roberts KL, Manicassamy B, & Lamb RA (2015) Influenza A virus uses intercellular connections to spread to neighboring cells. J Virol 89(3):1537–1549.

[26] Rustom A, Saffrich R, Markovic I, Walther P, & Gerdes HH (2004) Nanotubular highways for intercellular organelle transport. Science 303(5660):1007–1010.

[27] Thayanithy V, Dickson EL, Steer C, Subramanian S, & Lou E (2014) Tumor-stromal cross talk: direct cell-to-cell transfer of oncogenic microRNAs via tunneling nanotubes. Transl Res. 164(5):359–365.

[28] Wang X & Gerdes HH (2012) Long-distance electrical coupling via tunneling nanotubes. Biochim Biophys Acta 1818(8):2082–2086.

[29] Wang X & Gerdes HH (2015) Transfer of mitochondria via tunneling nanotubes rescues apoptotic PC12 cells. Cell Death Differ 22(7):1181–1191.

[30] Wang X, Veruki ML, Bukoreshtliev NV, Hartveit E, & Gerdes HH (2010) Animal cells connected by nanotubes can be electrically coupled through interposed gap-junction channels. Proc Natl Acad Sci USA 107(40):17194–17199.

[31] Wang X, et al. (2015) Rescue of Brain Function Using Tunneling Nanotubes Between Neural Stem Cells and Brain Microvascular Endothelial Cells. Mol Neurobiol 53(4):2480–2488.

[32] Wang Y, Cui J, Sun X, & Zhang Y (2011) Tunneling-nanotube development in astrocytes depends on p53 activation. Cell Death Differ 18(4):732–742.

[33] Zhu S, Victoria GS, Marzo L, Ghosh R, & Zurzolo C (2015) Prion aggregates transfer through tunneling nanotubes in endocytic vesicles. Prion 9(2):125–135.

[34] Mueller F, et al. (2013) FISH-quant: automatic counting of transcripts in 3D FISH images. Nat Methods 10(4):277–278.

[35] Huttelmaier S, et al. (2005) Spatial regulation of beta-actin translation by Src-dependent phosphorylation of ZBP1. Nature 438(7067):512–515.

[36] Katz ZB, et al. (2012) beta-Actin mRNA compartmentalization enhances focal adhesion stability and directs cell migration. Genes Dev 26(17):1885–1890.

[37] Buxbaum AR, Wu B, & Singer RH (2014) Single beta-actin mRNA detection in neurons reveals a mechanism for regulating its translatability. Science 343(6169):419–422.

[38] Yoon YJ, et al. (2016) Glutamate-induced RNA localization and translation in neurons. Proc Natl Acad Sci USA 113(44):E6877–e6886.

[39] Cabili MN, et al. (2015) Localization and abundance analysis of human lncRNAs at single-cell and single-molecule resolution. Genome Biol 16(1):20.

[40] Yunger S, Rosenfeld L, Garini Y, & Shav-Tal Y (2010) Single-allele analysis of transcription kinetics in living mammalian cells. Nat Methods 7(8):631–633.

[41] Palchaudhuri R, et al. (2015) A Small Molecule that Induces Intrinsic Pathway Apoptosis with Unparalleled Speed. Cell Rep 13(9):2027–2036.

[42] Tang EH & Vanhoutte PM (2008) Gap junction inhibitors reduce endothelium-dependent contractions in the aorta of spontaneously hypertensive rats. J Pharmacol Exp Ther 327(1):148–153.

[43] Sowinski S, Alakoskela JM, Jolly C, & Davis DM (2011) Optimized methods for imaging membrane nanotubes between T cells and trafficking of HIV-1. Methods 53(1):27–33.

[44] Schiller C, et al. (2013) LST1 promotes the assembly of a molecular machinery responsible for tunneling nanotube formation. J Cell Sci 126(Pt 3):767–777.

[45] Wu B, Chao JA, & Singer RH (2012) Fluorescence fluctuation spectroscopy enables quantitative imaging of single mRNAs in living cells. Biophys J 102(12):2936–2944.

[46] Davis DM & Sowinski S (2008) Membrane nanotubes: dynamic long-distance connections between animal cells. Nat Rev Mol Cell Biol 9(6):431–436.

[47] Moffitt JR, et al. (2016) High-throughput single-cell gene-expression profiling with multiplexed error-robust fluorescence in situ hybridization. Proc Natl Acad Sci USA 113(39):11046–11051.

[48] Svensson V, et al. (2017) Power analysis of single-cell RNA-sequencing experiments. Nat Methods. EPub Mar 6, 2017.

[49] Haimovich G, Choder M, Singer RH, & Trcek T (2013) The fate of the messenger is pre-determined: A new model for regulation of gene expression. Biochim Biophys Acta 1829(6-7):643–653.

[50] Buxbaum AR, Haimovich G, & Singer RH (2015) In the right place at the right time: visualizing and understanding mRNA localization. Nat Rev Mol Cell Biol 16(2):95–109.

[51] Park HY, et al. (2014) Visualization of dynamics of single endogenous mRNA labeled in live mouse. Science 343(6169):422–424.

[52] Austefjord MW, Gerdes HH, & Wang X (2014) Tunneling nanotubes: Diversity in morphology and structure. Commun Integr Biol 7(1):e27934.

[53] Hase K, et al. (2009) M-Sec promotes membrane nanotube formation by interacting with Ral and the exocyst complex. Nat Cell Biol 11(12):1427–1432.

[54] Ohno H, Hase K, & Kimura S (2010) M-Sec: Emerging secrets of tunneling nanotube formation. Commun Integr Biol 3(3):231–233.

[55] Ropars V, et al. (2016) The myosin X motor is optimized for movement on actin bundles. Nat Commun 7:12456.

[56] Milo R. PR (2016) Cell Biology by the Numbers (Garland Science, Taylor & Francis Group LLC, New York, NY, USA).

[57] McCaffrey MW & Lindsay AJ (2012) Roles for myosin Va in RNA transport and turnover. Biochem Soc Trans 40(6):1416–1420.

[58] Sotelo JR, et al. (2013) Myosin-Va-dependent cell-to-cell transfer of RNA from Schwann cells to axons. PloS One 8(4):e61905.

[59] Pasquier J, et al. (2013) Preferential transfer of mitochondria from endothelial to cancer cells through tunneling nanotubes modulates chemoresistance. J Transl Med 11:94.

[60] Rogers RS & Bhattacharya J (2013) When cells become organelle donors. Physiology (Bethesda) 28(6):414–422.

[61] Haimovich G, Cohen-Zontag O, & Gerst JE (2016) A role for mRNA trafficking and localized translation in peroxisome biogenesis and function? Biochim Biophys Acta 1863(5):911–921.

[62] Kraut-Cohen J & Gerst JE (2010) Addressing mRNAs to the ER: cis sequences act up! Trends Biochem Sci 35(8):459–469.

[63] Lesnik C, Golani-Armon A, & Arava Y (2015) Localized translation near the mitochondrial outer membrane: An update. RNA Biol 12(8):801–809.

[64] Weis BL, Schleiff E, & Zerges W (2013) Protein targeting to subcellular organelles via MRNA localization. Biochim Biophys Acta 1833(2):260–273.

[65] Lou E, et al. (2012) Tunneling nanotubes provide a unique conduit for intercellular transfer of cellular contents in human malignant pleural mesothelioma. PloS One 7(3):e33093.

[66] Sancho M, et al. (2014) Altered mitochondria morphology and cell metabolism in Apaf1-deficient cells. PloS One 9(1):e84666.

[67] Padovan-Merhar O, et al. (2015) Single mammalian cells compensate for differences in cellular volume and DNA copy number through independent global transcriptional mechanisms. Mol Cell 58(2):339–352.

[68] Sanders TA, Llagostera E, & Barna M (2013) Specialized filopodia direct long-range transport of SHH during vertebrate tissue patterning. Nature 497(7451):628–632.

[69] Follenzi A & Naldini L (2002) Generation of HIV-1 derived lentiviral vectors. Methods Enzymol 346:454–465.

[70] Lasser C, Eldh M, & Lotvall J (2012) Isolation and characterization of RNA-containing exosomes. J Vis Exp (59):e3037.

[71] Zenklusen D & Singer RH (2010) Analyzing mRNA expression using single mRNA resolution fluorescent in situ hybridization. Methods Enzymol 470:641–659.

[72] Schindelin J, et al. (2012) Fiji: an open-source platform for biological-image analysis. Nat Methods 9(7):676–682.

[73] Rajlabimagetools (https://bitbucket.org/arjunrajlaboratory/rajlabimagetools/wiki/Home.

